# Multiple metabolic signals including AMPK and PKA regulate glucose-stimulated double strand break resection in yeast

**DOI:** 10.1101/2021.02.26.433101

**Authors:** Stephanie Lomonaco, Dominic Bazzano, Thomas E. Wilson

## Abstract

DNA double strand breaks (DSBs) are cytotoxic lesions repaired by non-homologous end joining (NHEJ) and homologous recombination (HR), with 5’ strand resection being the committed step in transition from NHEJ to HR. We previously discovered that *gal1* yeast, which cannot metabolize galactose, were unable to perform efficient 5’ resection even though DSBs were formed. Adding glucose or restoring *GAL1* restored resection, suggesting that carbon source metabolism signals to DSB repair. Here we demonstrate that any fermentable carbon source, including raffinose, can stimulate resection and that the stimulatory effect of glucose was associated with decreased, not increased, cellular ATP. The effect was cell cycle dependent and did not occur in G1, while glucose augmented the G2/M checkpoint arrest even in cells deficient in resection. AMP-activated protein kinase pathway mutants showed only low basal resection despite glucose addition but had normal checkpoint arrest, indicating a primary role for Snf1 specifically in glucose-stimulated resection. The metabolic inputs to resection were multifactorial, however, with loss of the transcriptional repressor Mig1 leading to increased basal resection, three distinct patterns of deficiency with loss of the protein kinase A catalytic subunits, Tpk1, Tpk2 andTpk3, and a resection delay in yeast lacking the lysine demethylase Rph1 that helped separate early and late phase responses to glucose. These results reveal multiple interrelated metabolic signals that optimize DSB resection efficiency while independently amplifying the G2/M checkpoint response.

## Introduction

Non-homologous end joining (NHEJ) and homologous recombination (HR) are the two predominant DNA double strand break (DSB) repair pathways in eukaryotic cells [1]. HR is initiated by resection of the 5′-terminated DSB strands to generate the 3′ single-stranded DNA (ssDNA) that is required for Rad51 binding and strand invasion and prevents further possibility for religation of the DSB ends via NHEJ. Therefore, resection initiation is a critical and highly regulated control point in the choice between HR and NHEJ [1]. Whereas NHEJ is used throughout the cell cycle, HR is activated in S and G2 phases when sister chromatids are available as repair templates through the actions of cyclin-dependent kinase (CDK) in stimulating resection [2]. In turn, the ssDNA generated by DSB resection acts as a signal leading to checkpoint arrest of the cell cycle at G2/M, although yeast cells can eventually adapt to the checkpoint and continue with cell division despite a persistent DSB [3].

In a prior study of point mutants of DNA ligase IV, the NHEJ ligase, we made a surprising observation that DSB resection was strongly impaired in *gal1* mutant yeast that were unable to ferment the galactose used to induce formation of the DSB by the HO endonuclease [4]. Restoration of the *GAL1* gene or addition of glucose strongly stimulated DSB resection as measured by low resolution Southern blot analysis. Because DSBs formed efficiently, which required extensive metabolic capacity to synthesize HO, this glucose-dependent stimulation of resection argued against a simple energetic effect and suggested that metabolic signaling might influence HR resection alongside the canonical DNA damage response.

Since our 2013 study, many studies have continued to reveal that DSB repair and especially HR can indeed be influenced by carbon metabolites and metabolic enzymes [5–8]. 2-hydroxyglutarate, an oncometabolite produced by the neomorphic activity of somatically mutated isocitrate dehydrogenase 1 and 2 (IDH1/2) enzymes, is associated with a functional HR deficiency secondary to the ability of 2-hydroxyglutarate to inhibit α-ketoglutarate-dependent enzymes such as the histone demethylase KDM4 ([9]. The cancer metabolites fumarate and succinate also compete with α-ketoglutarate for binding to KDM4B resulting in reduced HR factor recruitment to DSBs [9, 10]. HR factors can also associate with metabolic enzymes. Leshets et al. showed that Sae2, an early component of HR and a positive resection factor, functionally interacts with fumarase, an enzyme that catalyzes the hydration of fumarate to L-malate [5].

The cellular response to glucose is controlled through two main pathways in yeast, the AMP-activated protein kinase (AMPK) Snf1 and protein kinase A (PKA). AMPK is a heterotrimeric complex that regulates glycolytic signaling [11]. Snf1 is phosphorylated during glucose deprivation by Sak1 and other kinases ([12–14] and helps regulate glucose levels and the expression of genes required for metabolism of alternative sugars through a feedback loop involving a protein complex formed with the glycolytic enzyme hexokinase, Hxk2, and the transcriptional repressor, Mig1 [15, 16]. Snf1 has been implicated in the DNA damage response through SUMOylation catalyzed by Mms21, a SUMO ligase [17], and its mammalian orthologue has been shown to regulate both the exonuclease Exo1 and NHEJ via the checkpoint protein 53BP1 [18, 19].

In yeast, PKA activity arises from three distinct heterotetrameric protein complexes that each have two regulatory subunits encoded by *BCY1* and two catalytic subunits encoded by one of three related genes, *TPK1*, *TPK2*, and *TPK3* [20]. cAMP regulates PKA activity by binding to Bcy1, alleviating its inhibitory effect on the catalytic subunits. The PKA pathway is activated by glucose, although this regulation can be complex. PKA activity can be active but reduced even in glucose media, which suggests high fermentative growth capacity rather than glucose availability as a main controlling factor of PKA activity *in* vivo [21]. PKA has also been shown to be involved in suppression of HR through phosphorylation of a BRCAness gene, EMSY [22], while yeast Tpk1 has been seen to reduce the efficiency of NHEJ [23].

Given the increasing intersection between metabolism and DNA repair, we used a tightly controlled yeast DSB system to explore the potential ways that carbon source and related metabolic signaling might lead to altered resection efficiencies. A precisely quantitative assay revealed multiple apparent contributors to net resection including basal resection in the absence of glucose and two apparent phases of stimulated resection after glucose addition. These studies, including yeast mutants in important carbon regulatory pathways, demonstrated that resection stimulation relies on a fermentable carbon source and is an active process dependent on full activation of Snf1. The phenomenon was cell cycle dependent in that glucose could not stimulate resection in G1, whereas G2/M arrest increased after glucose addition independently of resection.

## Results

### Fermentable carbon sources increase DSB resection efficiency

The site-specific DSB system we employed makes use of an HO endonuclease cut site placed into a nucleosome-free region of the *ILV1* promoter (Supplementary Figure 1) [24]. HO is expressed from the native *GAL1* locus such that it is induced by galactose addition while rendering the cells *gal1* mutant and unable to metabolize galactose from the first step, galactokinase. Most strains used in this study carried several additional mutations of relevance. A *bar1* mutation promoted efficient G1 synchronization by α-factor. A previously described point mutation, *dnl4*-K466A, abrogated activity of the yeast NHEJ ligase, Dnl4, while still supporting its proper recruitment to DSBs [4]; yeast were thus unable to repair the HO-induced DSB by NHEJ such that resection was obligatorily progressive (Figure 1A). A *MATa*-inc allele protected the yeast mating type locus from digestion by HO.

**Figure 1.**
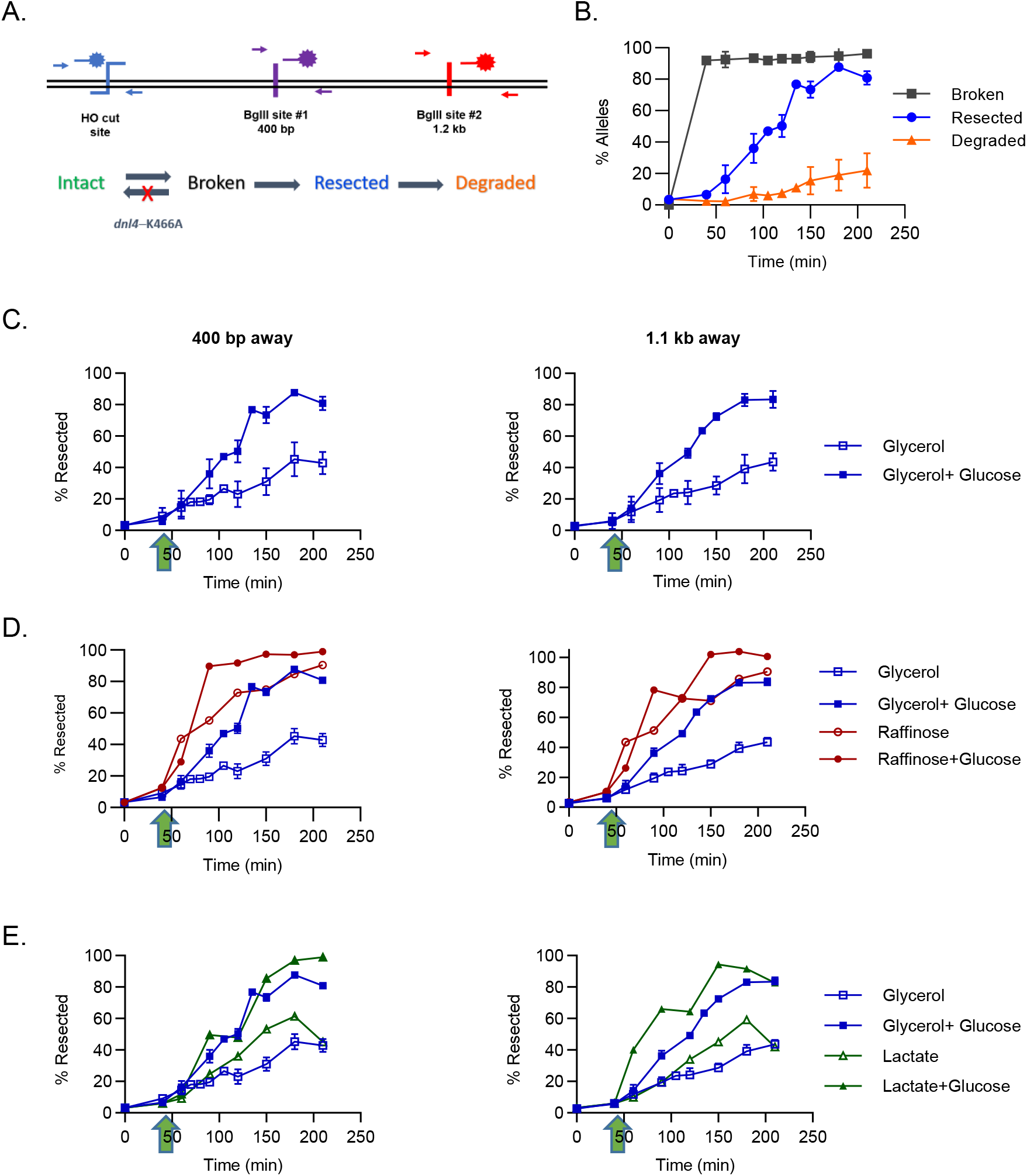
Fermentable carbon sources stimulate DSB resection. **(A)** ddPCR assay sites in the *ILV1* locus and progression through different states of DSB repair, where a *dnl4*-K466A mutation in most strains prevented NHEJ-mediated state reversal from broken back to intact. **(B)** Example ddPCR assay results showing quantification of the three measured states of broken, resected and degraded molecules. Note that the plotted resected stated is cumulative and thus includes molecules in the degraded state (see Methods); only the cumulative resected value is shown on subsequent plots. **(C)** Resected molecule fractions at 400 bp and 1.2 kb away from the HO cut site after DSB induction in glycerol, with and without glucose addition at 45 minutes. **(D)** Similar to (C) after pre-growth in raffinose. **(E)** Similar to (C) after pre-growth in lactate. Results are the mean +/− standard deviation of three biological replicates.

We measured DSB repair and resection using ddPCR for highest quantitative accuracy. Briefly, one set of primers flanked the HO cut site to detect DSB formation, one set resided in the distant *ACT1* gene to act as a control for comparing DNA amount between PCR tubes, and a final set flanked a BglII restriction enzyme site either 400 or 1200 bp from the HO cut site (Figure 1A). Adding BglII to DNA extracts only cut unresected dsDNA molecules whereas ssDNA resulting from 5’ resection persisted and gave signal [25] that we express as a percentage of resected molecules among all DSB-containing alleles (see Methods). Continued degradation of the 3’-terminated DSB strand results in complete loss of signal even without BglII addition (Figure 1B).

We first sought to revisit our prior finding with the improved resection assay by inducing DSBs by adding galactose to cells growing in glycerol. DSBs formed reliably and quickly within 30-40 minutes (Figure 1C). Unlike the apparent absence of resection in our prior Southern blots [4], with ddPCR we observed a measurable and steady progression of resection onset over time in cells maintained in glycerol + galactose (Figure 1C). Nevertheless, the resection rate increased markedly upon the addition of glucose 45 minutes after DSB induction (Figure 1C). We also noted what appeared to be a subtly biphasic response to glucose with a still further increase after 120 minutes (see more below).

To determine if the observed resection stimulation was specific to glucose, we further tested a series of fermentable and non-fermentable carbon sources. We pre-grew our yeast strain in the non-repressing carbon sources raffinose, glycerol, or lactate overnight and then added galactose to induce DSBs (Supplementary Figure 1). Notably, resection in raffinose cultures, which is fermentable but non-repressing, initiated very soon after DSB formation and very efficiently even without glucose addition, even though glucose was able to stimulate resection still further (Figure 1D). In contrast, yeast pre-grown in either glycerol or lactate, which are non-repressing and non-fermentable, showed lower basal levels of resection with sharp increases after glucose addition (Figure 1E). The biphasic response to glucose was especially pronounced in lactate (Figure 1E). Taken together, these data show that a fermentable carbon source, preferentially glucose, increases the efficiency of DSB resection.

### Glucose-stimulated resection is dependent on cell cycle stage

Because carbon source can stimulate changes in cell cycle distribution and resection is increased in S/G2 [26], we next compared results from asynchronous cultures (as above) to yeast synchronized in G1 or G2/M by treatment with α-factor or nocodazole, respectively. G1 arrest was highly efficient but G2/M arrest in glycerol cultures was incomplete (Supplementary Figure 2). We observed that cells arrested in G1 did not undergo a glucose-stimulated resection, but that the basal level of resection was the same as asynchronous cultures (Figure 2A). In contrast, nocodazole-treated cells behaved very similarly to asynchronous cultures, including both the same basal level of resection and a potent glucose stimulatory effect (Figure 2A). This pattern demonstrates that basal resection was independent of cell cycle stage, but that glucose-stimulated resection only occurred outside of G1.

**Figure 2.**
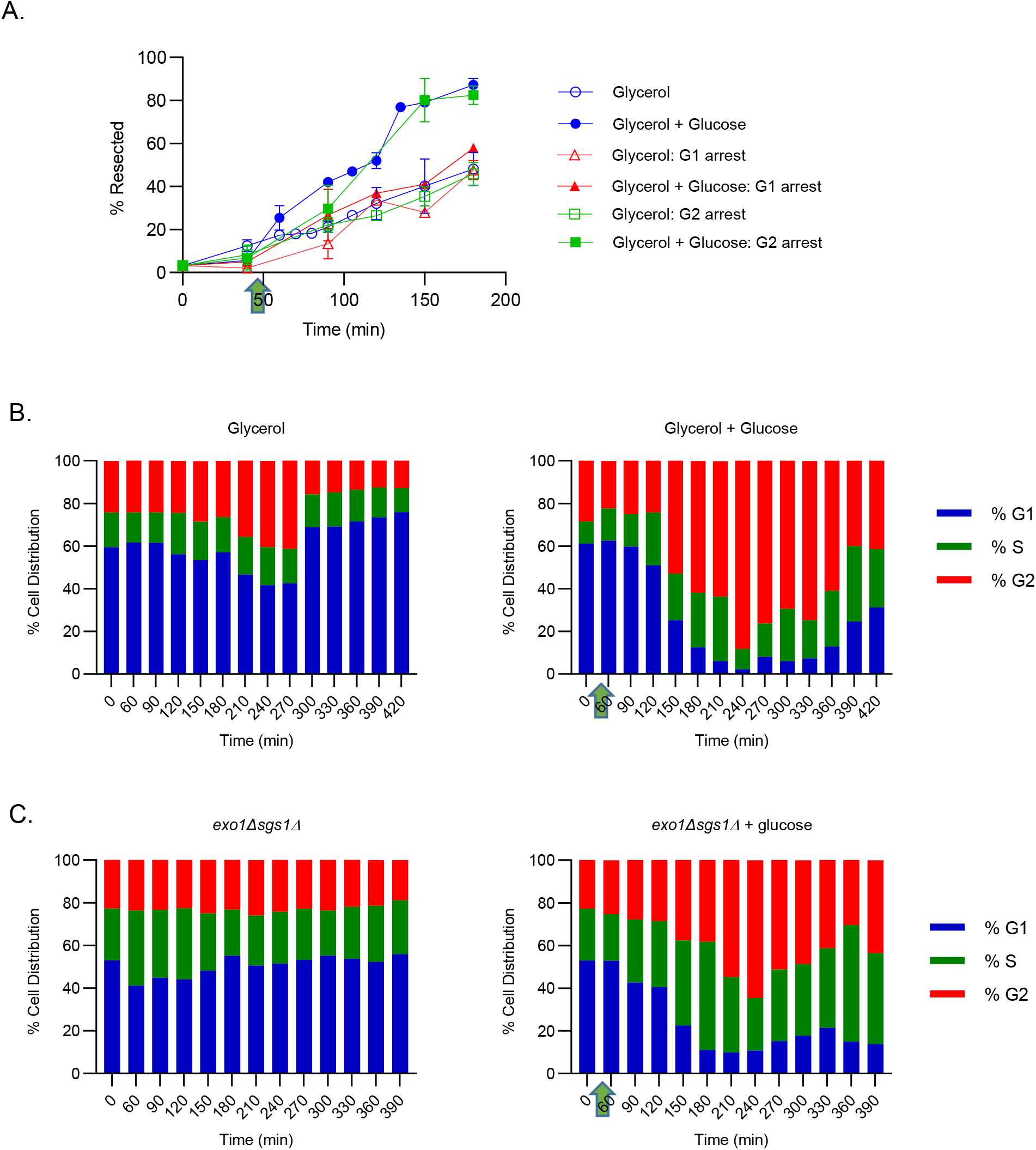
Glucose stimulates DSB resection specifically in S or G2/M and potentiates the G2/M checkpoint. **(A)** Resection efficiency in G1 and G2 arrested cells as compared to asynchronous cultures. **(B)** Cell cycle distributions of wild-type cells as determined by DNA content after DSB induction at time 0, with and without glucose addition at 45 minutes (arrow). Checkpoint arrest is evident as an increase in the %G2 and adaption as a subsequent decrease in %G2 from the peak value. **(C)** Similar to (B) for *exo1 sgs1* double mutant cells with substantially impaired resection.

### Glucose addition potentiates G2/M checkpoint arrest

In the converse of the previous experiments, we next examined the effect of DSB formation and altered resection rates on the cell cycle distribution, including the propensity to arrest or adapt to the DSB-dependent checkpoint at G2/M [3] as determined by DNA content over time. Cells held in glycerol + galactose showed a slight accumulation of G2/M cells but adapted to the G2/M checkpoint between 270 and 300 minutes (Figure 2B). In contrast, cells treated with glucose showed a much more potent and prolonged G2 arrest, consistent with the idea that the ssDNA resulting from higher levels of resection plays a role in G2 arrest (Figure 2B) [3]. To determine if that was the case we added both *exo1* and *sgs1* mutations to our strain background to nearly abolish resection [27]. This strain showed no alteration in the cell cycle distribution in response to DSB formation in glycerol + galactose (Figure 2C). Interestingly, we did see a measurable G2/M arrest and adaptation response in *exo1 sgs1* mutant cells after glucose addition, although the double mutant only accumulated to 40% G2/M-phase cells at 240 minutes as opposed to 80% for wild-type cells (Figure 2C). These results suggest that there is more than simply extensive ssDNA formation that is causing G2 arrest and that glucose addition potentiates the checkpoint arrest signal independently of extensive resection.

### Metabolic changes in response to carbon source and DSBs

To further explore how glucose might be stimulating resection we investigated the metabolite profile of cells through the time course of our DSB induction paradigm using metabolomics (see Methods). To distinguish the metabolic impact of carbon source changes themselves from the DSB formation and resection induced by those changes, we used two strains with or without a competent HO cut site in the *ILV1* gene promoter. Cells without the cut site (no DSB) reveal only carbon source dependent changes while those with the cut site can reveal how the DSB might alter that metabolic response. We further compared three different culture trajectories. The first was maintained in glycerol and sampled at 0 and 150 minutes to serve as a baseline against which other effects could be compared. The second two were both exposed to galactose at 0 minutes, with one further subjected to glucose addition at 45 minutes.

Figure 3A summarizes the results from a targeted metabolite panel focused on glycolysis, the TCA cycle and nucleotide metabolites [28] (complete data are provided in Supplementary Files 1 and 2). As expected, glucose addition generally increased the levels of intermediates in early glycolysis and the pentose phosphate pathway. Glucose also markedly increased cellular lactate levels (LAC), consistent with our inference that fermentation is a key state associated with increased resection. The early TCA intermediates citrate (CIT) and succinate (SUC) also increased, but the later intermediates fumarate (FUM) and malate (MAL) decreased, although individually these were not always significant changes (potential implications are discussed further below). Interestingly, even the addition of unfermentable galactose depressed cellular ATP levels (Supplementary File 1), which glucose addition drove still lower while also increasing AMP levels (Figure 3A). As a result, the energy charge was the lowest after glucose addition even though this is when resection activity was the highest (Figure 3B), which we attribute to the cells entering a consumptive anabolic phase (also discussed further below). Finally, we noted that the presence of the DSB typically did not change these metabolite responses, although the 3-carbon intermediates glycerol 3-phosphate (Gl-OH-3P) and 2-phosphoglycerate/3-phosphoglycerate (2PG/3PG) were increased by glucose only when the DSB was present. We do not know the basis of this effect and it will not be discussed further.

**Figure 3.**
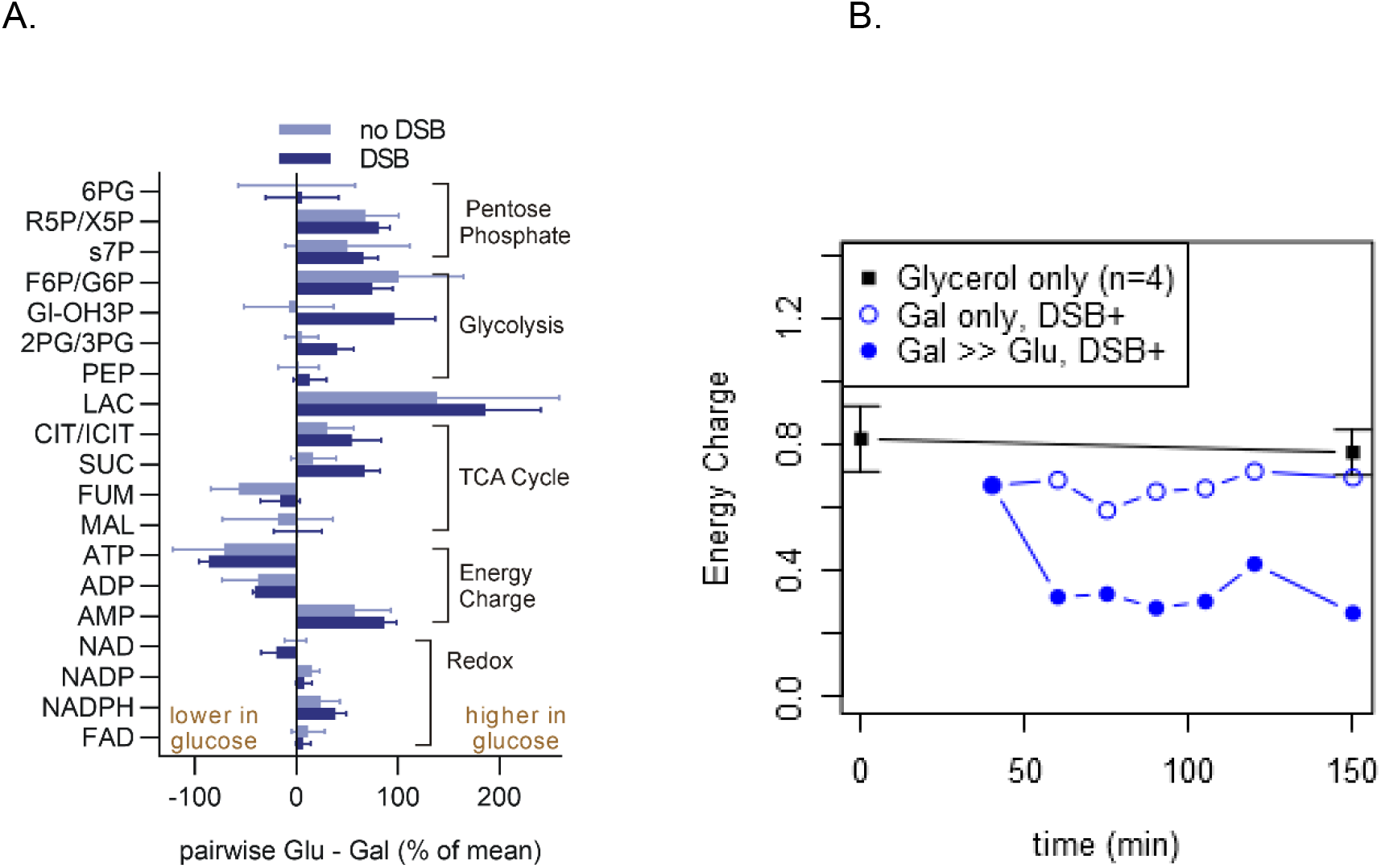
Glucose increases glycolytic and early TCA intermediates while suppressing late TCA intermediates and cellular energy charge, independently of the DSB. **(A)** Summary of most metabolites in the Gly-TCA targeted metabolomics panel grouped by pathway. For each metabolite, two values are plotted corresponding to strains with a competent (DSB) or incompetent (no DSB) HO cut site at the *ILV1* locus. Plotted values represent the extent to which glucose increased or decreased the metabolite level (see Methods for details), as the average +/− standard error of 5 measurements from 60 to 120 minutes. **(B)** Expanded time course of the energy charge calculated from the ATP, ADP and AMP concentrations. The black line represents four independent measurements at the beginning and end of the time course for cultures treated with neither galactose nor glucose. When indicated, glucose was added at 45 minutes.

### The Snf1 pathway plays a critical role in glucose-stimulated resection

Armed with a description of resection stimulation by fermentable carbon sources and its relationship with cell cycle and metabolism, we next sought to identify metabolic signals that could mediate these effects by making targeted mutations informed by the patterns above. We first explored an axis suggested by the results with adenylate intermediates. Specifically, Snf1 is the yeast AMPK [29] and thus might be expected to be responsive to the changing energy charge noted in Figure 3, although glucose might prevent this activity through the activation of Reg1/Glc7 and subsequent dephosphorylation of the Snf1 activation loop [11]. For these and other reasons we sought to explore a potential role of Snf1 in the resection response.

Yeast lacking Snf1 do not grow in glycerol media, so we instead deleted Snf1 complex subunits important for its activation and translocation to the nucleus. Sak1 is an AMPK kinase that phosphorylates Snf1 and allows Snf1 to bind to the Gal83 subunit and translocate to the nucleus [30, 31]. Strikingly, deleting either *SAK1* or *GAL83* abrogated glucose-stimulated resection while having no impact on basal resection levels in glycerol + galactose (Figure 4A). There also was a large cell cycle arrest at G2 in the *sak1*Δ strain after glucose addition (Figure 4B) as well as in the *gal83*Δ mutant strain (not shown), with limited adaptation at later time points. These results strongly implicate Snf1 activity as being required for glucose stimulation of resection but not a normal checkpoint response.

**Figure 4.**
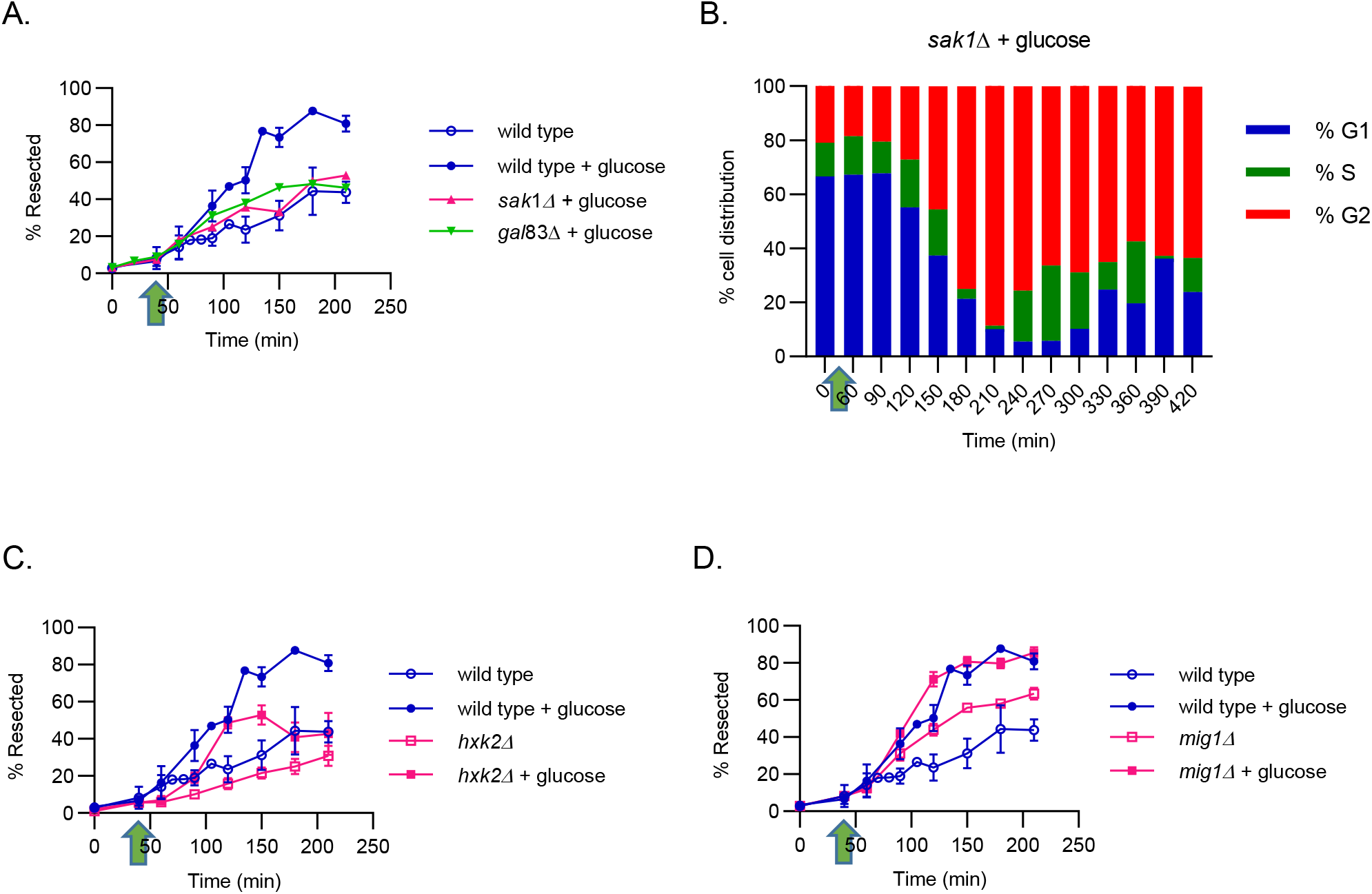
Mutations of Snf1-associated glucose-sensing pathways alter the DSB resection response. **(A)** Resection efficiency 400 bp away from the HO DSB in strains bearing deletion mutations in genes encoding Snf1 complex subunits, *SAK1* and *GAL83*. Plots and time courses are similar to Figure 1. **(B)** Cell cycle distributions of *sak1*Δ cells as determined by DNA content after DSB induction at time 0, with and without glucose addition at 45 minutes (arrow). **(C)** Similar to (A) for a *mig1* deletion strain. **(D)** Similar to (A) for a *hxk2* deletion strain. Results are the mean +/− standard deviation of three biological replicates.

To examine this axis further we also mutated the genes encoding Hxk2 and Mig1. The hexokinase, Hxk2, plays two distinct roles in the cell as a glycolytic enzyme in the cytosol and as a regulator of transcription in the nucleus [32]. Specifically, Hxk2 is a regulator of glucose repression through its nuclear localization and structural association with the transcriptional repressor Mig1 [16, 33]. Snf1 binds to Hxk2 and allows for phosphorylation and subsequent release of Mig1 and export to the cytoplasm, releasing the repressive ability of Mig1 [15].

Deleting *HXK2* resulted in a distinctive pattern wherein *hxk2* mutant yeast showed a reduced basal level of resection and at least a 30-minute delayed in glucose stimulated resection (Figure 4C). Finally, deleting *MIG1* resulted in yet a third distinctive pattern with an increased basal level of resection and retention of a normal profile in glucose (Figure 4D). These changes in the initial resection response as seen with *hxk2* mutant suggest an immediate glucose-stimulated response through metabolic signaling and a second response through transcriptional regulation, as seen by derepression of glucose regulation in the *mig1* mutant.

### Rph1, a demethylase, plays an early role in glucose-stimulated resection

The modest depression of fumarate by glucose addition in Figure 3A was interesting because the TCA cycle metabolites fumarate and succinate have been shown to have a negative impact on HR efficiency [5, 6, 8, 9]. Our data are consistent with a model in which glucose might stimulate resection by alleviating that inhibitory effect by decreasing fumarate levels. In human cells, fumarate and other oncometabolites inhibit HR by a mechanism in which those metabolites inhibit the histone lysine demethylase KDM4B [9, 34]. Rph1 is the yeast homolog of KDM4 [35] so we mutated it next. As seen in Figure 5A, *rph1* mutant yeast showed yet another distinctive resection pattern where basal resection was unaffected and glucose did stimulate resection but only in what appeared to correspond to the second of the two phases of the normally biphasic response. An alternative view is that *rph1* mutation delayed the normal response to glucose. This suggests that an early component, such as DNA accessibility to the break (e.g. through chromatin modification) may play a role in glucose-stimulated resection.

**Figure 5.**
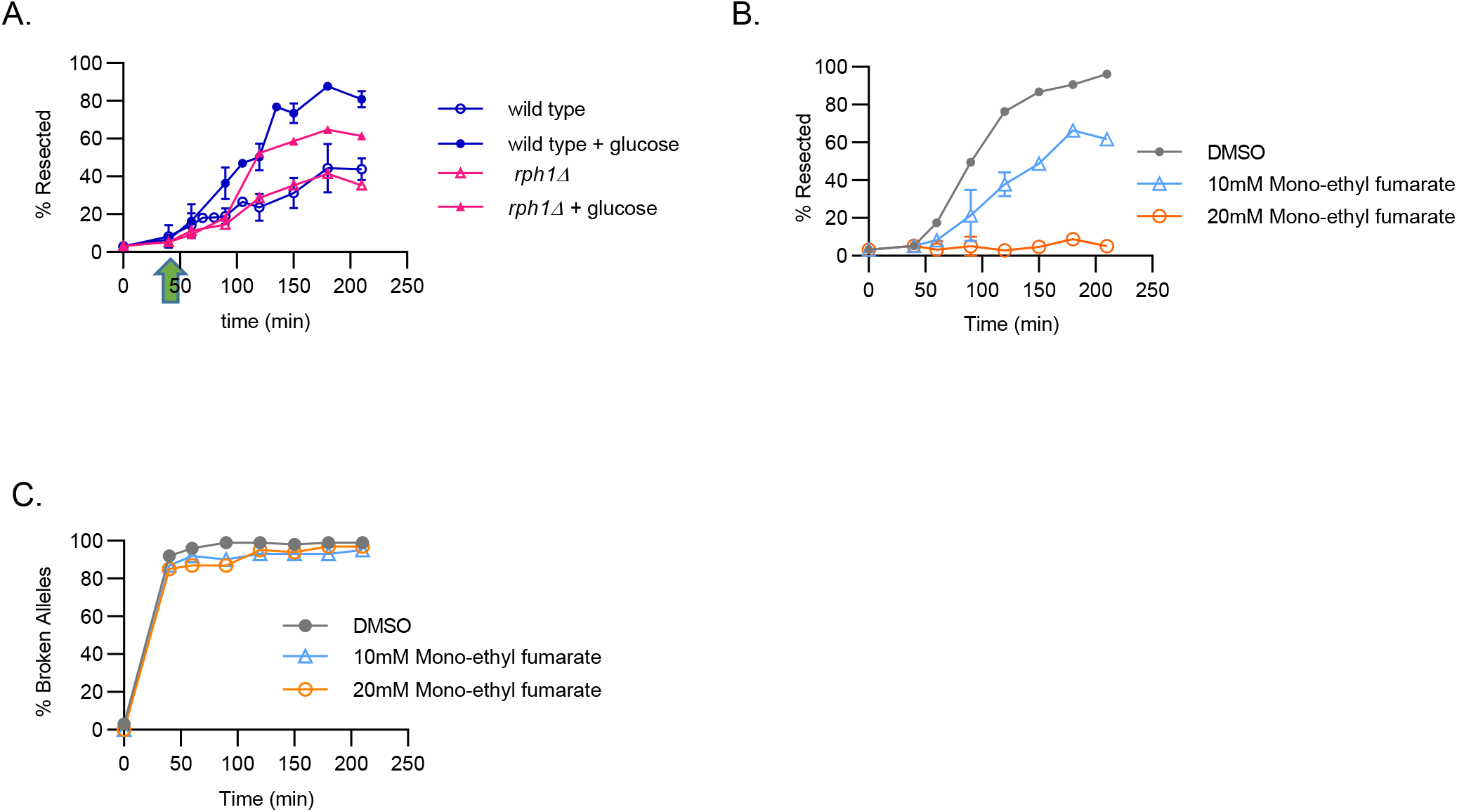
The Rph1 demethylase plays a role in early glucose-stimulated resection that is not entirely explained by fumarate. **(A)** Resection efficiency 400 bp away from the HO DSB in a *rph1* mutant strain. Plots and time courses are similar to Figure 1. **(B)** Resection efficiency similar to (A) for the MEF-treated cultures. **(C)** Fraction of broken *ILV1* alleles after 1 hour of pre-incubation in mono-ethyl fumarate (MEF) at the indicated concentrations followed by induction of the DSB by galactose at 0 minutes and glucose addition at 45 minutes. MEF pre-incubation did not prevent the DSB from forming. Results are the mean +/− standard deviation of three biological replicates.

The results above were not fully consistent with the fumarate hypothesis in that cells with the *rph1* mutation were able to resect more extensively than cells without glucose addition. To examine this in more detail we added varying amounts of monomethyl fumarate (MEF) to cultures before and during DSB formation and resection to artificially increase cellular fumarate levels. Similar to reports by others [5], fumarate at sufficient concentrations was remarkably able to completely suppress all DSB resection (Figure 5B). Importantly, cells did not die or become inactive despite pre-growth in MEF because the DSB was formed normally (Figure 5C), an outcome which requires considerable cellular activity to synthesize the HO enzyme.

### PKA subunits play unique roles in glucose-stimulated resection

As stated above PKA has a complex role in cellular glucose responses with its primary role as a nutrient sensing complex. There has also been recent evidence to suggest overlap of the glucose regulating pathways mediated by Snf1 and PKA [36, 37], as well as evidence that PKA subunits play a role in DSB repair [38]. We also see this complexity in the resection response of PKA mutant strains (Figure 6) wherein loss of the three different catalytic subunits of different PKA complexes all showed different resection patterns. Deleting *TPK1* mainly led to a reduction of basal resection while retaining a substantial glucose-stimulated resection response above that low baseline level, albeit to a lesser net resection efficiency than wild-type cells in glucose (Figure 6A). Deleting *TPK2* had a more subtle phenotype but appeared to selectively impair only the latter of the two phases of the normal biphasic response to glucose (Figure 6B). Finally, deleting *TPK3* increased the basal level of resection (Figure 6C).

**Figure 6.**
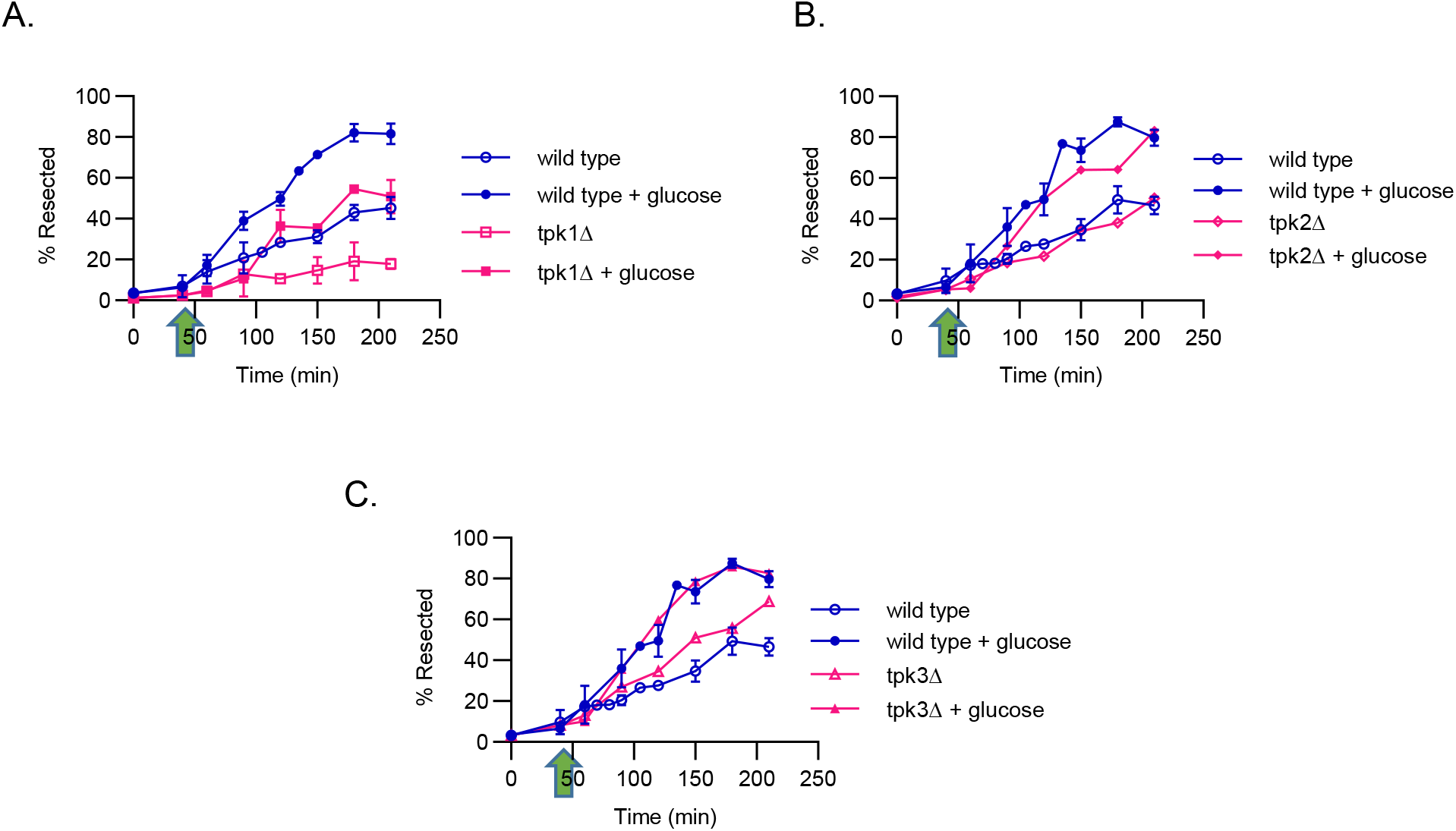
PKA catalytic subunits all play unique roles in glucose-stimulated resection. Resection efficiency 400 bp away from the HO DSB in strains bearing deletion mutations in the genes encoding the PKA catalytic subunits **(A)** *TPK1*, **(B)** *TPK2* and **(C)** *TPK3*. Plots and time courses are similar to Figure 1. Results are the mean +/− standard deviation of three biological replicates.

## Discussion

Results in this study demonstrate that yeast DSB resection is strongly influenced by carbon sources and their metabolism and that these effects are mediated by a surprisingly complex set of distinct and separable influences (Figure 7). The basic result was that adding glucose to glycerol cultures markedly increases DSB resection efficiency. Our prior finding that galactose could only stimulate resection in *GAL1* wild-type yeast that could use it as a carbon source suggested that the ability to ferment a sugar was the key parameter associated with increased resection. Indeed, studies here of other carbon sources established that raffinose, which is fermentable, confers rapid resection while lactate, which like glycerol is non-fermentable, does not. Metabolomics analysis confirmed that glucose increased early glycolytic intermediates and lactate, consistent with the transition to fermentation.

**Figure 7.**
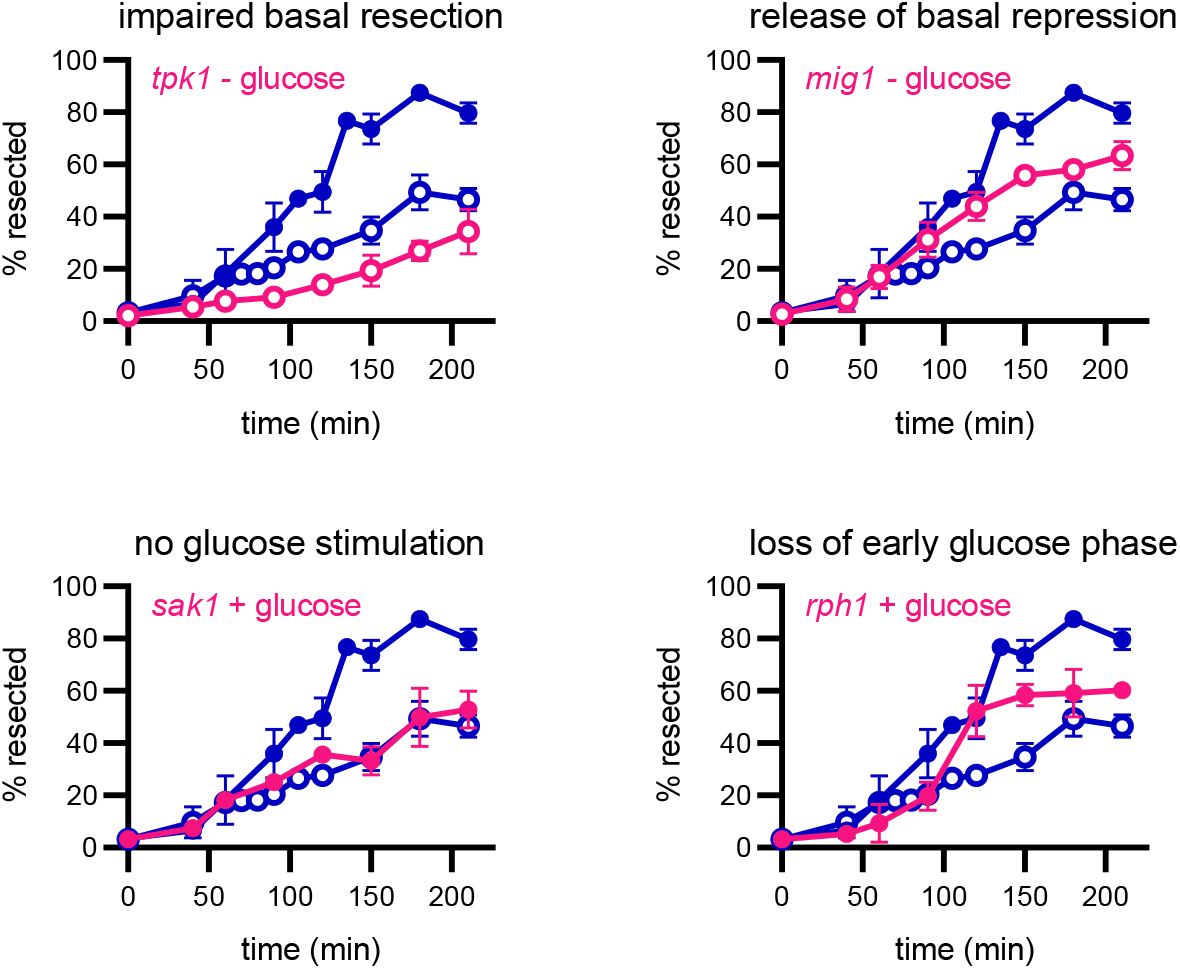
Multiple patterns of resection efficiency dependent on metabolic signaling reveal multiple layers of resection control. Data panels show only selected traces to illustrate distinctive resection patterns dependent on carbon metabolism (magenta) as compared to the wild-type pattern of basal plus glucose-stimulated resection repeated in all panels (blue). Note that some mutants (magenta) are shown with, some without, glucose addition. See text for further discussion.

We were previously concerned that increased resection in glucose might simply have been due to lower available energy in glycerol. Metabolomics showed this was not the case because cells with the highest rates of resection in glucose had the lowest ATP levels and energy charge and highest AMP levels. Here, it is noteworthy that base removal in resection does not require energy, although other actions such as chromatin displacement do. Other published evidence further supports the idea that glucose metabolism is important in resection progression and DSB repair independently of simply providing more energy for resection, including that decreased ATP leads to increased DNA processing in the DNA damage response [39]. Cancer cells have aberrant glucose metabolism and demonstrate resistance to many chemotherapeutic agents because of enhanced DNA repair. The aberrant glucose metabolism may increase nucleotide pools that could increase DSB repair efficiency [40]. However, after addition of glucose to our system, our data do not indicate an increased nucleotide pool, even with increased resection. They instead suggest that metabolism of glucose is itself a major contributor to resection efficiency.

We were also concerned that the beneficial role of glucose in resection might be solely do to the facts that glucose stimulates the transition of cells from G1 to S via PKA-mediated signaling [41] and that CDK activation in S phase leads to increased resection via covalent modifications of pro-resection proteins such as Sae2 [42]. Several lines of reasoning indicate this is not the case and that the impacts of carbon source are substantially more complex. It is true that glucose was unable to stimulate resection in G1. However, cells with a substantially increased G2/M fraction due to treatment with nocodazole showed the same pattern as asynchronous cells and all wild-type cultures had the same basal level of resection without glucose addition. If cells in S/G2 always had higher resection due to cell cycle signals we would have expected cell cycle stage manipulation alone to have largely recapitulated the glucose effect and to blunt the effect of glucose in G2/M, neither of which occurred. Mutants discussed below further distinguish the glucose and cell cycle effects.

Cell cycle progression studies provided further evidence for a distinct action of glucose signaling in G2/M phase cells. Haber and colleagues showed that yeast follow a sequence in which persistent DSBs, such as the one in this study, lead to checkpoint arrest of the cell cycle that is stimulated by resected ssDNA, but that over 12 hours cells adapt to the checkpoint and divide even with the unrepaired DSB [3]. We saw the same phenomenon in our cells but noted that the degree of checkpoint arrest was lesser without glucose, which is again inconsistent with the idea that all cells in G2 behave the same. Moreover, *exo1 sgs1* double mutant cells with a severe resection defect still showed a notable increase in checkpoint arrest in response to glucose. Finally, *sak1* mutant yeast did not stimulate resection in response to glucose but still had a potent glucose-stimulated checkpoint arrest. Together, we suggest that the stimulatory effect of glucose is potentiated by a crosstalk with cell cycle signals [43, 44] such that the glucose effect can only be realized in S/G2 but that it is nevertheless an independent signal from the cell cycle with separable influences on resection and the G2/M checkpoint.

Our further work investigating metabolic signaling mutants was most revealing of the multiple ways that glucose and other fermentable carbon sources stimulate resection. If the effect was simple, we might have expected a small number of mutant phenotypes but instead observed many as summarized in Figure 7. There was a basal amount of resection irrespective of cell cycle stage, but that resection could be lowered by mutations including *hxk2* and *tpk1*. Basal resection could also be selectively increased in *mig1* yeast, supporting the argument that both positive and negative factors contribute to resection observed in G1 or without glucose. Additionally, glucose stimulated resection in what appears to be a biphasic manner, with an early and a late response phase. This pattern was very subtle and difficult to defend based on wild-type data alone, but two mutants, *rph1* and *tpk2* appeared to impair glucose stimulation in the early and late phases, respectively. While these exact interpretations require further investigation, the different patterns are most easily explained by the existence of multiple metabolic inputs to resection efficiency.

The most striking result was that loss of Sak1 and Gal83 essentially abrogated the glucose stimulatory effect on resection. These mutations inhibit Snf1 translocation to the nucleus in glycerol [45], indicating that nuclear localization of Snf1 and by inference Snf1 catalytic activity are important for glucose to stimulate resection. Snf1 also binds to Hxk2 in the nucleus and allows for phosphorylation and subsequent release of Mig1 and its export to the cytoplasm, releasing the transcription repression conferred by Mig1 [15]. We saw distinct effects of the loss of Hxk2 and Mig1 consistent with this pattern wherein Hxk2 protein promoted while Mig1 protein repressed basal resection. These different results have several paradoxes that require further investigation. First, Snf1 is generally activated and required in low glucose yet promotes resection in response to glucose. The resolution to that paradox may lie in the priming of cells to respond to glucose due to prior activation of Snf1. Second, Mig1 acts as a transcriptional repressor in glycerol yet represses resection in glucose. Here, the resolution may lie in a non-transcriptional role of Mig1 or alternatively that Mig1 represses expression of a factor inhibitory to resection.

Recent papers have shown that the lysine demethylase KDM4B is inhibited by high levels of fumarate and other oncometabolites, which causes aberrant hypermethylation of histones decreasing proper recognition of the DSB [9, 10]. Also, the enzyme fumarase directly contributes to resection by an interaction with the pro-resection protein Sae2 [5]. Our metabolomics profiles suggested that glucose lowered cellular fumarate levels and offered an additional potential mechanism for the glucose effect on resection. Indeed, yeast lacking the yeast KDM4 homologue Rph1 showed decreased glucose-stimulated resection at early time points but a robust delayed, i.e. later phase, response to glucose. These results help to establish the idea of a biphasic response noted above but further work will be required to determine if the effect is secondary to H3K36 methylation of histones, given that yeast lack the H3K9 methyl mark identified as most important of the KDM4B effect in human cells [46]. We further note that fumarate cannot be the entire basis of the mechanism for glucose stimulation due to Rph1 retaining a significant response. Nevertheless, fumarate proved to be an exceptionally potent resection inhibitor when added exogenously as MEF.

Finally, our data suggest complex resection regulation dependent on the catalytic subunits of the glucose-sensing PKA pathway. We see that resection is inhibited in glucose in a *tpk1*Δ strain. The *tpk2*Δ mutant strain showed a delayed yet robust resection efficiency compared to wild type while the *tpk3*Δ mutant strain had an increased basal resection. Previous studies have shown that the three TPK genes are positively regulated by the Snf1 signaling pathway when glycerol and glucose are utilized as the carbon source [36]. However, the PKA promoters are all differentially regulated in response to carbon source. Further studies are needed to distinguish whether Snf1 and Tpk1 functional interactions are involved in regulation of DSB repair.

In summary, our results support a model in which yeast DSB resection comprises a basal activity under both positive and negative regulation that can be potently stimulated in a biphasic response to fermentable carbon sources outside of G1. This multiplicity of phenotypic patterns showcases a complex crosstalk between glucose sensing pathways, the cell cycle and DNA repair. Results indicate that both DNA repair and damage checkpoints are independently impacted by glucose in a manner that cannot be explained by either cell cycle or energetic effects alone. Instead, the phenotypes of relevant mutant strains suggest that altered protein complex formation, transcriptional regulation and chromatin modifications might all be active contributors to fine tuning resection efficiency based on the cellular environmental context.

## Materials and Methods

### Yeast strains and growth media

Haploid yeast used in this study were isogenic derivatives of BY4741 [47]. The base strain for most strains used in this study was YW3104, itself a derivative of YW2566 that has been previously described [24]. The construction of YW3104 is described in detail in Bazzano et al (in preparation). Additional gene disruptions and modified alleles were made using a polymerase chain reaction (PCR)-mediated technique or a *URA3* pop-in/pop-out method and confirmed by PCR and sequencing [48]. A table of all yeast strains is provided as Supplementary Table 1.

Yeast were grown at 30°C in either rich medium containing 1% yeast extract, 2% peptone and 40 μg/ml adenine (YPA) with either 2% glucose, 2% galactose, 2% glycerol, 2% lactate, or 2% raffinose as a carbon source. When indicated, α-factor (Sigma-Aldrich T6901) was added at a final concentration of 5 μg/ml and nocodazole (Sigma-Aldrich M1404) was added from a stock at 1 mg/ml in DMSO to a final concentration of 7.5 μg/ml. Mono-ethyl fumarate (MEF, Sigma-Aldrich 128422) was added from a stock at 200 mM in DMSO to a final concentration of 10 or 20 mM. Equivalent amounts of DMSO alone were added to control cultures when studying the effects of nocodazole or MEF.

### DSB induction paradigm

To ensure maximal consistency in DSB induction profiles, yeast were first grown overnight in YPA-dextrose and then diluted back into YPA-glycerol to an OD_600_ of 0.02. Cultures were incubated overnight with shaking at 250 RPM to an OD_600_ in the range 0.4 to 0.6 to ensure logarithmic growth. When indicated, cells were synchronized by addition of α-factor or nocodazole or pre-treated with MEF at this stage of handling. Galactose was then added to the media to a final concentration of 2% to induce the DSB followed by continued shaking at 30C. Cultures without glucose addition were maintained in this way, while cells treated with glucose were pelleted at 3000 RPM for 5 min and resuspended in YPA-dextrose media. Samples of cultures were harvested at indicated time points, pelleted and flash frozen with dry ice, then stored at −80C until needed.

### Double strand break and resection assay by ddPCR

Genomic DNA was extracted from frozen cell pellets using the ThermoFisher Yeast DNA Extraction Kit and resuspended in TE. Genomic DNA (400 ng) was either mock digested (i.e. without enzyme addition) or digested with BglII (New England Biolabs, 5 units/rxn, NEBuffer 3.1) for 1 hour at 37°C. Samples were then diluted 1:5 and placed into a 96 well plate with Supermix (no dUTPs, BioRad) and the required target locus primers and MGB-NFQ probes (Applied Biosystems; see Supplementary Table 2). Three tubes were set up for each source DNA time point. All contained a control probe (VIC) targeting the non-DSB *ACT1* locus to report on the exact amount of DNA added per tube. Tube #1 was BglII-undigested and additionally had a FAM probe with primers flanking the HO cut site (the DSB tube). Tube #2 was BglII-undigested and additionally had a FAM probe with primers flanking the BglII site at the resection query position (the undegraded DNA tube). Tube #3 was BglII-digested and additionally had a FAM probe with primers flanking the BglII site at the resection query position (the resected tube). ddPCR was executed in a BioRad QX200 AutoDG Droplet Digital PCR System. PCR cycling conditions were 95 C for 10 min, 40 cycles: 94 C for 30 sec, 60 C for 1 min, final 98 C for 10 min, hold at 4 C. Data were analyzed using the vendor-provided QuantaSoft software to yield final amplifiable molecule concentrations in each tube in copies per μl.

Further calculations were performed from the final probe-derived molecule concentrations from each of the three tubes as follows. First, we established primary values for the DNA states that we directly measured. The <u>total</u> amount of cellular equivalents of DNA was established from the control locus probe concentration in Tube #1 (referred to as ‘total’ in calculations below, etc.). The amount of <u>intact</u> (i.e. uncut) DSB sites was established by the HO probe concentration from Tube #1. The amount of <u>undegraded</u> DSB DNA at the resection query position was established by the BglII probe concentration from mock-digested Tube #2. The amount of <u>resected</u> DSB DNA (prior to correction for BglII cleavage efficiency) at the resection query position was established by the BglII probe concentration from BglII-digested Tube #3.

We next applied needed data corrections. First, to correct for any errors due to pipetting, the undegraded and resected values from Tubes #2 and #3 were normalized based on the *ACT1* control probe concentrations from the three tubes to place them on the same numeric scale as the values from Tube #1. Second, we accounted for incomplete BglII digestion using the time 0 sample by calculating the <u>fractionUndigested</u> DNA as ‘resected_t=0_ / undegraded_t=0_’, given that at time 0 no DSB had formed and thus no DNA could have been truly resected. Corrected resected values accounting for this digestion efficiency error were calculated as ‘(resected – undegraded * fractionUndigested) / (1 – fractionUndigested)’, where the logic was to remove from the resected value the amount of undegraded DNA that is inferred to be undigested. The undigested fraction was assumed to apply equally to all samples. Because digestion efficiency was consistently high, this correction was typically small.

Next we calculated secondary values derived from the corrected primary values. The amount of <u>degraded</u> DSB DNA at the resection query position was calculated as ‘total – undegraded’. The amount of <u>broken</u> HO loci, i.e. DNA that had suffered the DSB, was calculated as ‘total – intact’. The final derived calculation accounted for the fact that once DSB DNA had passed from the resected (i.e. loss of only the 5’ terminated strand) to the degraded (i.e. loss of both strands) state it could no longer give signal as ‘resected’. We assumed that degradation always followed true 5’ resection. Because we wished to plot all molecules that had been resected up to time T, including those had continued on to complete degradation, our final plotted value for cumulative resected DNA was calculated as ‘resected + degraded’. Our last transformations simply expressed the final values of broken, resected and degraded DNA as a % of the total molecules. We provide an Excel spreadsheet as Supplementary File 3 that executes all of the above calculations for a single time series of measurements.

All experiments were repeated over at least two independent biological replicates. Final calculations aggregated data from those replicates, with values on plots expressed as the average +/− standard deviation.

### Flow cytometry

Yeast cultures were collected at various time points after DSB induction similar to the DSB resection assay above. Samples were fixed in 70% ethanol and stored at −20C. Samples were stained with propidium Iodide (Thermo-Fisher P1304MP) and subjected to flow on an ACEA NovoCyte flow cytometer in the Michigan Pathology Flow Cytometry Core. Data were analyzed using FlowJo 10 software to determine the fraction of cells in each of the G1, S and G2/M cell cycle stages utilizing the Watson cell cycle model.

### Metabolomics assessment

Yeast were grown in an analogous fashion to the DSB resection paradigm above. Samples were harvested (3 × 10^6^ cells), washed with 150 mM ammonium acetate at the indicated time points followed by dry pellet rapid freeze in liquid nitrogen and storage at −80C. Samples were transferred to the University of Michigan Metabolomics Core facility who performed targeted metabolite analysis using the Gly/TCA/nucleotide panel [28]. Briefly, LCMS detection included a one-step liquid-liquid organic solvent extraction cells and separation on a 1mm x150mm HILIC specific column in a 35 min cycle. All analytes were measured by electron spray ionization on a LC-QTOF mass spectrometer. When available, peak areas were normalized to internal standards to account for matrix effects and compared to a standard curve to determine final μM concentrations, which were finally converted to pmol per μg protein to allow sample comparisons.

Supplementary File 1 provides a summary of all metabolite names as well as detailed plots of each individual metabolite. To aggregate data to visualize the impact of glucose addition relative to cultures held in glycerol + galactose, we extracted the five data points from 60 to 120 minutes, the times showing the greatest changes in resection due to glucose stimulation. For each time point we calculated the difference between the galactose-only and glucose cultures as ‘glucose – galactose’. We took the mean and standard deviation of these pairwise differences and normalized them to a common scale by dividing them by the mean of all values for the target time points over both the galactose-only and glucose cultures. We calculated p-values for the difference between the carbon sources using a two-sided paired t-test. The energy charge was calculated as ‘([ATP] + [ADP] / 2) / (ATP + ADP + AMP)’. The R script that performed all of these calculations is provided as Supplementary File 4 and the source metabolomics data tables in Excel format in Supplementary File 2.

### Fumarate treatment

Yeast cultures after overnight growth in glycerol as described above were pre-treated with MEF at varying concentrations for 1 hour prior to the induction of DSBs by the addition of 2% galactose. Samples were transitioned to glucose media as described above. After 1 hour of continued shaking to allow either resection or repair to occur.

## Supporting information

Supplemental File 1 - metabolomics profiles

Supplemental File 2 - metabolomics data

Supplemental File 3 - ddPCR calculations

Supplemental File 4 - metabolomics script

## Acknowledgements

This work was supported by NIH grant R01 GM120767 to T.E.W and a supplement of the same number awarded to S.L. The authors thank Maureen Kachman of the Michigan Metabolomics Core for assistance and helpful discussions.

## Conflicts

The authors declare that they have no competing financial interest with this work.

## Notes

### Competing Interest Statement

The authors have declared no competing interest.

## References

1. Symington, L.S., Mechanism and regulation of DNA end resection in eukaryotes. Crit Rev Biochem Mol Biol, 2016. 51(3): p. 195–212.

2. Mao, Z., et al., DNA repair by nonhomologous end joining and homologous recombination during cell cycle in human cells. Cell Cycle, 2008. 7(18): p. 2902–6.

3. Lee, S.E., et al., Saccharomyces Ku70, mre11/rad50 and RPA proteins regulate adaptation to G2/M arrest after DNA damage. Cell, 1998. 94(3): p. 399–409.

4. Chiruvella, K.K., et al., Saccharomyces cerevisiae DNA ligase IV supports imprecise end joining independently of its catalytic activity. PLoS Genet, 2013. 9(6): p. e1003599.

5. Leshets, M., et al., Fumarase is involved in DNA double-strand break resection through a functional interaction with Sae2. Curr Genet, 2018. 64(3): p. 697–712.

6. Leshets, M., et al., Fumarase: From the TCA Cycle to DNA Damage Response and Tumor Suppression. Front Mol Biosci, 2018. 5: p. 68.

7. Sizemore, S.T., et al., Pyruvate kinase M2 regulates homologous recombination-mediated DNA double-strand break repair. Cell Res, 2018. 28(11): p. 1090–1102.

8. Ui, A., et al., Possible involvement of LKB1-AMPK signaling in non-homologous end joining. Oncogene, 2014. 33(13): p. 1640–8.

9. Sulkowski, P.L., et al., Oncometabolites suppress DNA repair by disrupting local chromatin signalling. Nature, 2020. 582(7813): p. 586–591.

10. Sulkowski, P.L., et al., Krebs-cycle-deficient hereditary cancer syndromes are defined by defects in homologous-recombination DNA repair. Nat Genet, 2018. 50(8): p. 1086–1092.

11. McCartney, R.R. and M.C. Schmidt, Regulation of Snf1 kinase. Activation requires phosphorylation of threonine 210 by an upstream kinase as well as a distinct step mediated by the Snf4 subunit. J Biol Chem, 2001. 276(39): p. 36460–6.

12. Hong, S.P., et al., Activation of yeast Snf1 and mammalian AMP-activated protein kinase by upstream kinases. Proc Natl Acad Sci U S A, 2003. 100(15): p. 8839–43.

13. Nath, N., R.R. McCartney, and M.C. Schmidt, Yeast Pak1 kinase associates with and activates Snf1. Mol Cell Biol, 2003. 23(11): p. 3909–17.

14. Sutherland, C.M., et al., Elm1p is one of three upstream kinases for the Saccharomyces cerevisiae SNF1 complex. Curr Biol, 2003. 13(15): p. 1299–305.

15. Ahuatzi, D., et al., The glucose-regulated nuclear localization of hexokinase 2 in Saccharomyces cerevisiae is Mig1-dependent. J Biol Chem, 2004. 279(14): p. 14440–6.

16. Fernandez-Garcia, P., et al., Phosphorylation of yeast hexokinase 2 regulates its nucleocytoplasmic shuttling. J Biol Chem, 2012. 287(50): p. 42151–64.

17. Kobi J Simpson-Lavy, A.B., Martin Kupiec, Mark Johnston, Cross-Talk between Carbon Metabolism and the DNA Damage Response in S. cerevisiae. Cell Reports, 2015. 12: p. 1865–1875.

18. Jiang, Y., et al., AMPK-mediated phosphorylation on 53BP1 promotes c-NHEJ. Cell Rep, 2021. 34(7): p. 108713.

19. Li, S., et al., Ca(2+)-Stimulated AMPK-Dependent Phosphorylation of Exo1 Protects Stressed Replication Forks from Aberrant Resection. Mol Cell, 2019. 74(6): p. 1123–1137 e6.

20. Portela, P. and S. Rossi, cAMP-PKA signal transduction specificity in Saccharomyces cerevisiae. Curr Genet, 2020. 66(6): p. 1093–1099.

21. Thevelein, J.M., et al., Nutrient-induced signal transduction through the protein kinase A pathway and its role in the control of metabolism, stress resistance, and growth in yeast. Enzyme Microb Technol, 2000. 26(9-10): p. 819–825.

22. Jelinic, P., et al., The EMSY threonine 207 phospho-site is required for EMSYdriven suppression of DNA damage repair. Oncotarget, 2017. 8(8): p. 13792–13804.

23. Hooshyar, M., et al., Deletion of yeast TPK1 reduces the efficiency of non-homologous end joining DNA repair. Biochem Biophys Res Commun, 2020. 533(4): p. 899–904.

24. Wu, D., L.M. Topper, and T.E. Wilson, Recruitment and dissociation of nonhomologous end joining proteins at a DNA double-strand break in Saccharomyces cerevisiae. Genetics, 2008. 178(3): p. 1237–49.

25. Zhou, Y. and T.T. Paull, Direct measurement of single-stranded DNA intermediates in mammalian cells by quantitative polymerase chain reaction. Anal Biochem, 2015. 479: p. 48–50.

26. Mathiasen, D.P. and M. Lisby, Cell cycle regulation of homologous recombination in Saccharomyces cerevisiae. FEMS Microbiol Rev, 2014. 38(2): p. 172–84.

27. Mimitou, E.P. and L.S. Symington, Sae2, Exo1 and Sgs1 collaborate in DNA double-strand break processing. Nature, 2008. 455(7214): p. 770–4.

28. Lorenz, M.A., C.F. Burant, and R.T. Kennedy, Reducing time and increasing sensitivity in sample preparation for adherent mammalian cell metabolomics. Anal Chem, 2011. 83(9): p. 3406–14.

29. Ye, T., et al., The mammalian AMP-activated protein kinase complex mediates glucose regulation of gene expression in the yeast Saccharomyces cerevisiae. FEBS Lett, 2014. 588(12): p. 2070–7.

30. Elbing, K., et al., Subunits of the Snf1 kinase heterotrimer show interdependence for association and activity. J Biol Chem, 2006. 281(36): p. 26170–80.

31. Hedbacker, K., S.P. Hong, and M. Carlson, Pak1 protein kinase regulates activation and nuclear localization of Snf1-Gal83 protein kinase. Mol Cell Biol, 2004. 24(18): p. 8255–63.

32. Pelaez, R., P. Herrero, and F. Moreno, Functional domains of yeast hexokinase 2. Biochem J, 2010. 432(1): p. 181–90.

33. Vega, M., et al., Hexokinase 2 Is an Intracellular Glucose Sensor of Yeast Cells That Maintains the Structure and Activity of Mig1 Protein Repressor Complex. J Biol Chem, 2016. 291(14): p. 7267–85.

34. Jiang, Y., et al., Local generation of fumarate promotes DNA repair through inhibition of histone H3 demethylation. Nat Cell Biol, 2015. 17(9): p. 1158–68.

35. Young, L.C., D.W. McDonald, and M.J. Hendzel, Kdm4b histone demethylase is a DNA damage response protein and confers a survival advantage following gamma-irradiation. J Biol Chem, 2013. 288(29): p. 21376–88.

36. Galello, F., et al., Transcriptional regulation of the protein kinase a subunits in Saccharomyces cerevisiae during fermentative growth. Yeast, 2017. 34(12): p. 495–508.

37. Kummel, A., et al., Differential glucose repression in common yeast strains in response to HXK2 deletion. FEMS Yeast Res, 2010. 10(3): p. 322–32.

38. Omidi, K., et al., Uncharacterized ORF HUR1 influences the efficiency of non-homologous end-joining repair in Saccharomyces cerevisiae. Gene, 2018. 639: p. 128–136.

39. Hopfner, K.P., ATP puts the brake on DNA double-strand break repair: a new study shows that ATP switches the Mre11-Rad50-Nbs1 repair factor between signaling and processing of DNA ends. Bioessays, 2014. 36(12): p. 1170–8.

40. Lin, J., et al., The roles of glucose metabolic reprogramming in chemo- and radio-resistance. J Exp Clin Cancer Res, 2019. 38(1): p. 218.

41. Searle, J.S., et al., The DNA damage checkpoint and PKA pathways converge on APC substrates and Cdc20 to regulate mitotic progression. Nat Cell Biol, 2004. 6(2): p. 138–45.

42. Matsuzaki, K., et al., Cyclin-dependent kinase-dependent phosphorylation of Lif1 and Sae2 controls imprecise nonhomologous end joining accompanied by double-strand break resection. Genes Cells, 2012. 17(6): p. 473–93.

43. Pessina, S., et al., Snf1/AMPK promotes S-phase entrance by controlling CLB5 transcription in budding yeast. Cell Cycle, 2010. 9(11): p. 2189–200.

44. Zhang, L., et al., Multiple Layers of Phospho-Regulation Coordinate Metabolism and the Cell Cycle in Budding Yeast. Front Cell Dev Biol, 2019. 7: p. 338.

45. Vincent, O., et al., Subcellular localization of the Snf1 kinase is regulated by specific beta subunits and a novel glucose signaling mechanism. Genes Dev, 2001. 15(9): p. 1104–14.

46. Klose, R.J., et al., Demethylation of histone H3K36 and H3K9 by Rph1: a vestige of an H3K9 methylation system in Saccharomyces cerevisiae? Mol Cell Biol, 2007. 27(11): p. 3951–61.

47. Brachmann, C.B., et al., Designer deletion strains derived from Saccharomyces cerevisiae S288C: a useful set of strains and plasmids for PCR-mediated gene disruption and other applications. Yeast, 1998. 14(2): p. 115–32.

48. Palmbos, P.L., J.M. Daley, and T.E. Wilson, Mutations of the Yku80 C terminus and Xrs2 FHA domain specifically block yeast nonhomologous end joining. Mol Cell Biol, 2005. 25(24): p. 10782–90.

