## Supplemental Figures for "Multiple metabolic signals including AMPK and PKA regulate glucose-stimulated double strand break resection in yeast"

**Supplementary Files**

**Supplementary File 1.** Composite view of all metabolomics profiles, including a table of targeted metabolites that passed quality control. See details in file.

**Supplementary File 2.** Excel file with primary metabolomics data. See details in file.

**Supplementary File 3.** Excel template for performing resection ddPCR calculations.

**Supplementary File 4.** R script that generated all metabolomics figures in the study. See comments in file.

**Supplementary Table 1: Yeast strains used in this work.**

| **Name** | **Genotype** |
|---|---|
| YW3104 | *MAT*a-inc::*LEU2* *can1*Δ::*ILV1*-QPCR *gal1*::HO *his3*Δ1 *ILV1*prm::HOcs *ILV1*Reg::Ori-His3MX6 *bar1*Δ *leu2*Δ0 *met15*Δ0 *ura3*Δ0 *dnl4*-K466A |
| YW3081 | YW3104 *mig1*Δ |
| YW3178 | YW3104 *sak1*Δ |
| YW3179 | YW3104 *gal83*Δ |
| YW3225 | YW3104 HOcs ACA deletion |
| YW3227 | YW3104 *rph1*Δ |
| YW3278 | YW3104 *tpk1*Δ |
| YW3279 | YW3104 *tpk2*Δ |
| YW3280 | YW3104 *tpk3*Δ |
| YW3284 | YW3104 *hxk2*Δ |

**Supplementary Table 2: Primers and probes for resection ddPCR assay.**

| **Name** | **Target** | **Sequence** |
|---|---|---|
| OW3058 | Forward primer *ACT1* | AGAGTTGCCCCAGAAGAACA |
| OW3059 | Reverse primer *ACT1* | GGCTTGGATGGAAACGTAGA |
| OW4100 | *ACT1* MGBLQ probe VIC | TGACTGAAGCTCCAATGAACCCT |
| OW3991 | Forward primer HOcs | AATAAGAAGGGCAAAAAGAAAAAGC |
| OW3992 | Reverse primer HOcs | AAAGCAGCAACAACAAAAGTTTTC |
| OW4221 | HO cut site MGBNFQ probe FAM | CGCTTTTAGTTTCAGCTTTCCGCA |
| OW4182 | Forward primer BglII 400 bp away | TCCCGTCTAAACACGAATGTCA |
| OW4183 | Reverse primer BglII 400 bp away | GGTGTACAAACAGGCATAACGATA |
| OW4184 | BglII site 400 bb MGNFQ probe FAM | ATCATGCCCAAGGTGTGGCC |
| OW4141 | Forward primer BglII 1.2 kb away | TGGTGAAGGAAAGGAAGTCT |
| OW4142 | Reverse primer BglII 1.2 kb away | GCTTGATTTCCTTTTCTCTGTCA |
| OW4140 | BglII site 1.2 kb MGBNFQ probe FAM | ACCCGACGTCCCTGGTGCGTTCA |

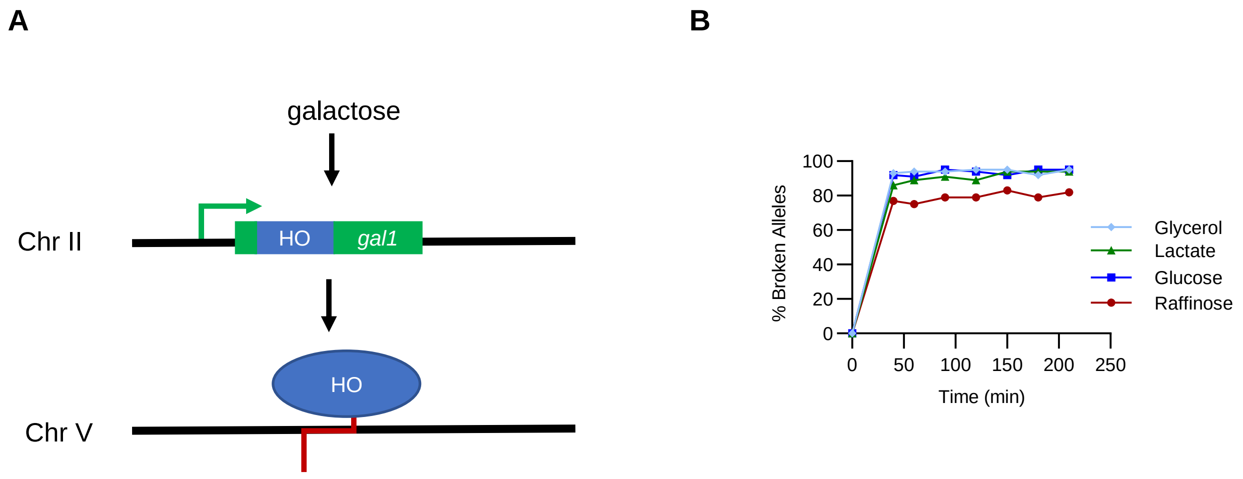


**Supplementary Figure 1. DSB induction systems. (A)** *gal1*-HO system, induced by galactose. **(B)** HO cutting efficiency of *gal1*-HO system in various carbon sources. The target HOcs and resulting DSB are in the native yeast *ILV1* promoter on ChrV.

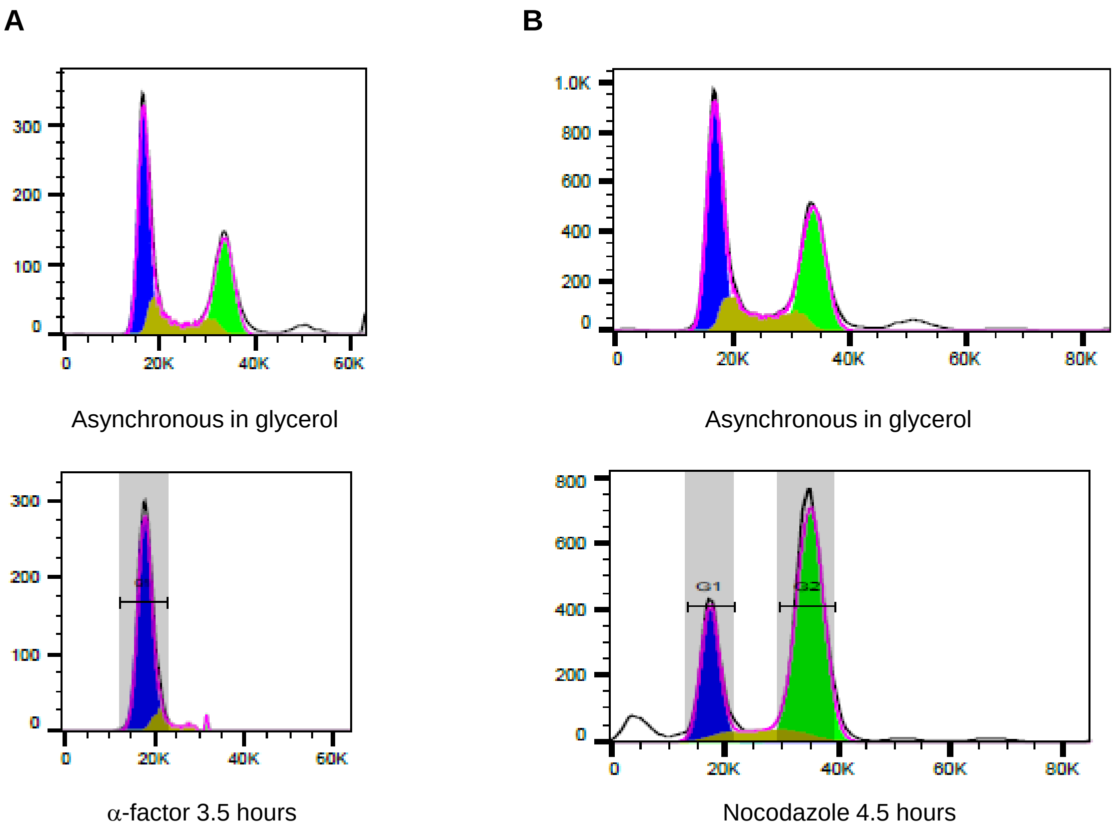


**Supplementary Figure 2. Flow cytometry DNA content histograms of G1 arrest and G2 arrest. (A)** Asynchronous culture in glycerol (top) and then after addition of α-factor (bottom). **(B)** Asynchronous culture in glycerol (top) and then after the addition of nocodazole (bottom). Cells growing in glycerol (unlike glucose) never achieved complete G2/M arrest in nocodazole but the fraction of G1 cells dropped considerably from 62% to 18%.
