## Supplemental File 1 - metabolomics profiles for "Multiple metabolic signals including AMPK and PKA regulate glucose-stimulated double strand break resection in yeast"

This file summarizes the metabolomics data sets. For each metabolite, two plots are drawn in units of pmol/μg total cellular protein.

The **left panel** shows a black line connecting the average +/- standard deviation of 4 independent samples measured at the 0 and 150 min time points (without any addition of glucose or galactose) as a reference point for the other traces. Blue and green lines represent cultures that did or did not have a competent HO cut site, respectively. Filled and open symbols indicate cultures that did or did not have 2% glucose added at 45 min, respectively.

The **right panel** shows the same data where the time points from 60 to 120 minutes are shown together for each of the four colored traces. A solid horizontal line is the average of all of the glycerol-only measurements. A dashed horizontal line is the average of all the data points on the plot.

Metabolites are listed in the following order. Some metabolites from the Gly-TCA panel could not be reported due to technical failures or values outside of the standard curve.

| Metabolite | Long Name | Category |
|---|---|---|
| **6PG** | 6-Phosphogluconate | pentose phosphate |
| **R5P/X5P** | Ribose 5-phosphate + Xylulose 5-phosphate | pentose phosphate |
| **s7P** | Sedheptulose 7-phosphate | pentose phosphate |
| **F6P/G6P** | Glucose-6-phospate + Fructose-6-Phosphate | glycolysis |
| **Gl-OH3P** | Glycerol 3-phosphate | glycolysis via DHAP |
| **2PG/3PG** | 3-Phosphoglycerate + 2-Phosphoglycerate | glycolysis |
| **PEP** | Phosphoenolpyruvate | glycolysis |
| **LAC** | Lactate | fermentation |
| **CIT/ICIT** | Citrate + Isocitrate | TCA cycle |
| **SUC** | Succinate | TCA cycle |
| **FUM** | Fumarate | TCA cycle |
| **MAL** | Malate | TCA cycle |
| **ATP** | ATP | adenylates |
| **ADP** | ADP | adenylates |
| **AMP** | AMP | adenylates |
| **NAD** | NAD | redox |
| **NADP** | NADP | redox |
| **NADPH** | NADPH | redox |
| **FAD** | FAD | redox |

### 6-Phosphogluconate

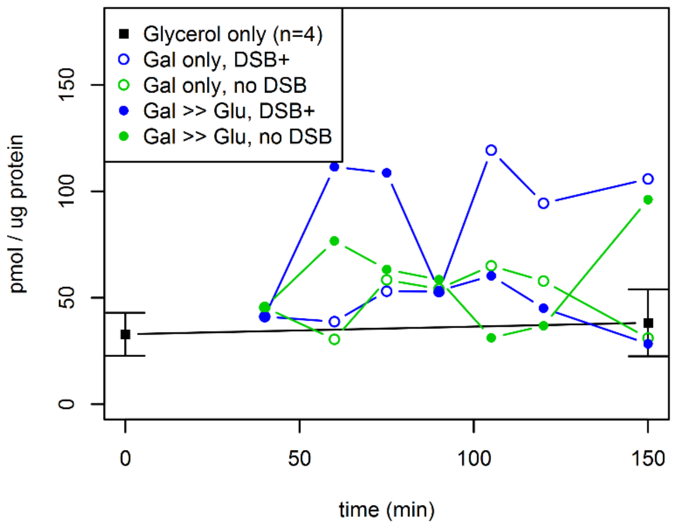


### 6-Phosphogluconate

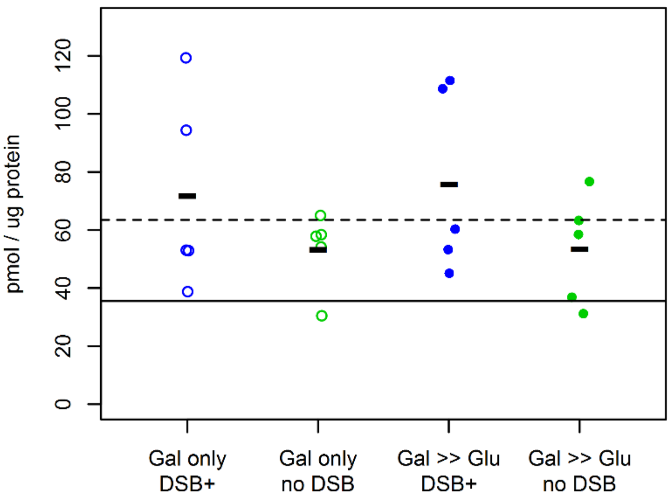


### Ribose 5-phosphate + Xylulose 5-phosphate

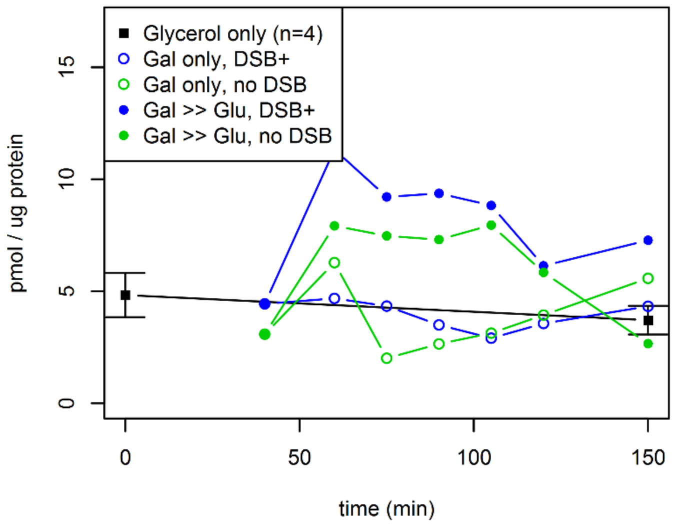


### Ribose 5-phosphate + Xylulose 5-phosphate

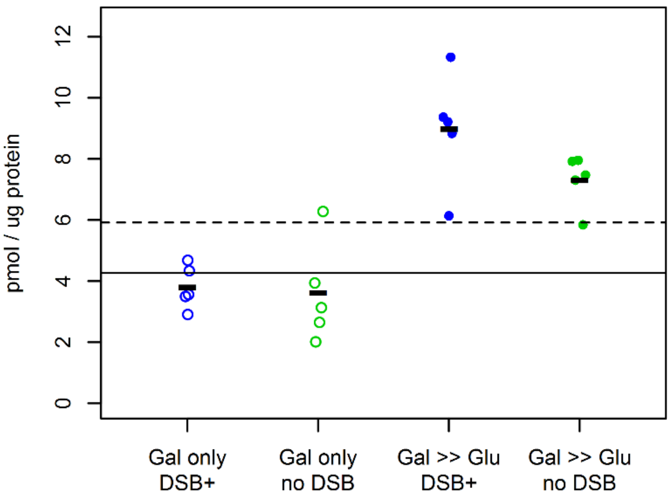


### Sedheptulose 7-phosphate

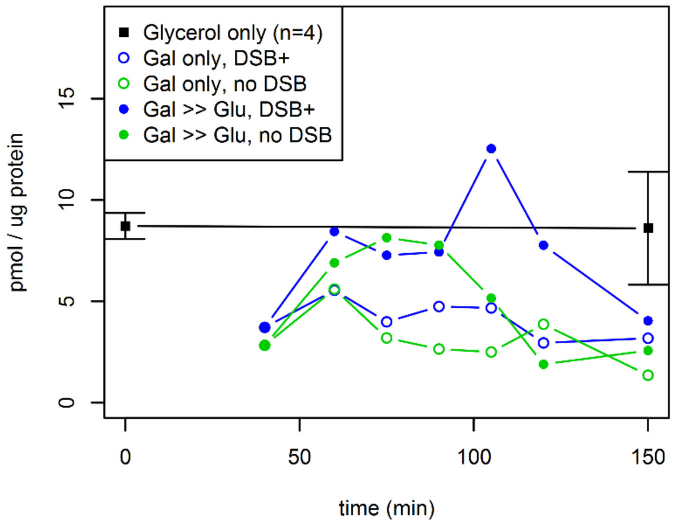


### Sedheptulose 7-phosphate

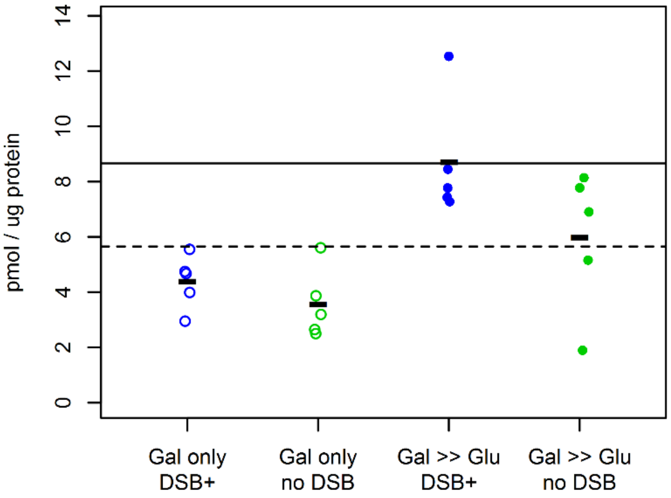

**Glucose-6-phospate + Fructose-6-Phosphate**

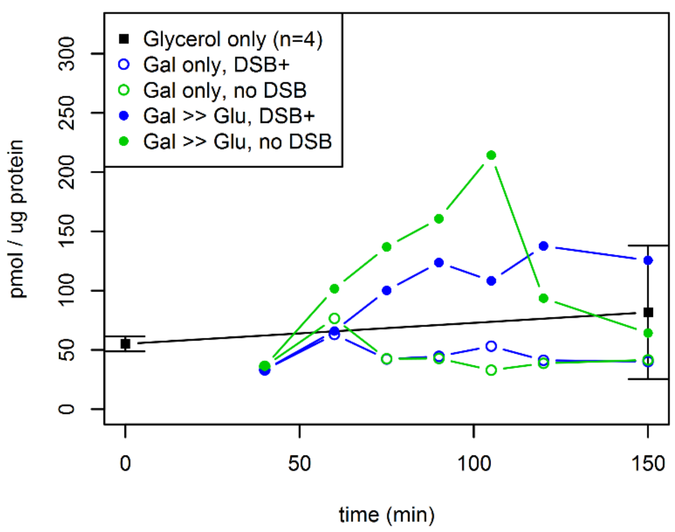


**Glucose-6-phospate + Fructose-6-Phosphate**

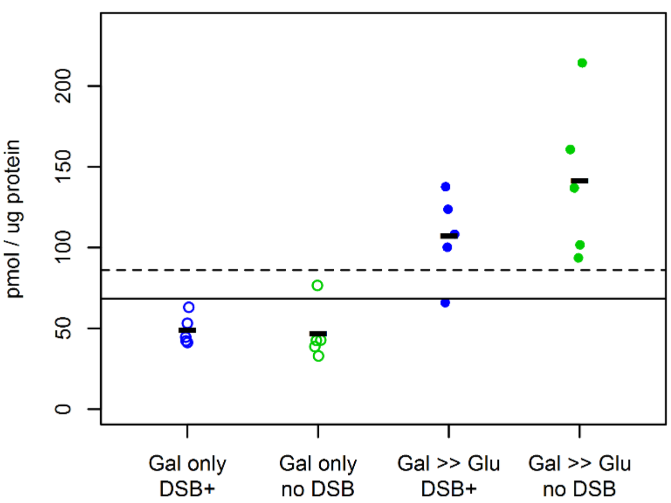


**Glycerol 3-phosphate**

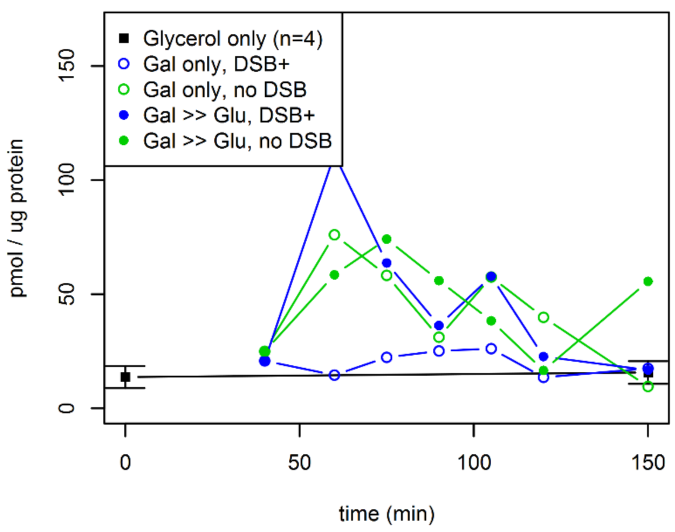


**Glycerol 3-phosphate**

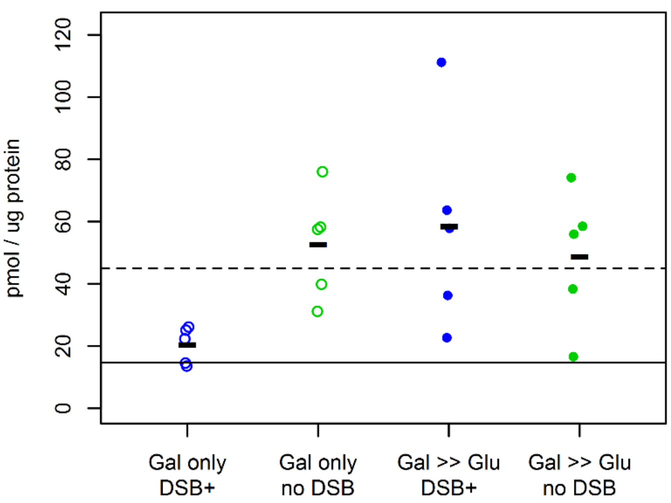


**3-Phosphoglycerate + 2-Phosphoglycerate**

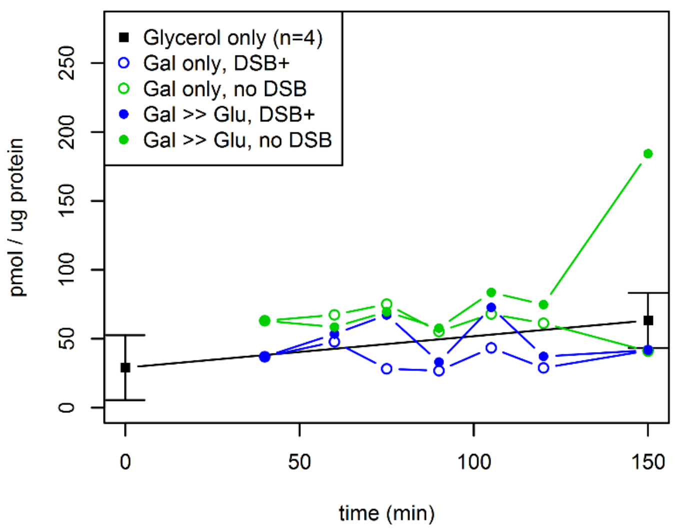


**3-Phosphoglycerate + 2-Phosphoglycerate**

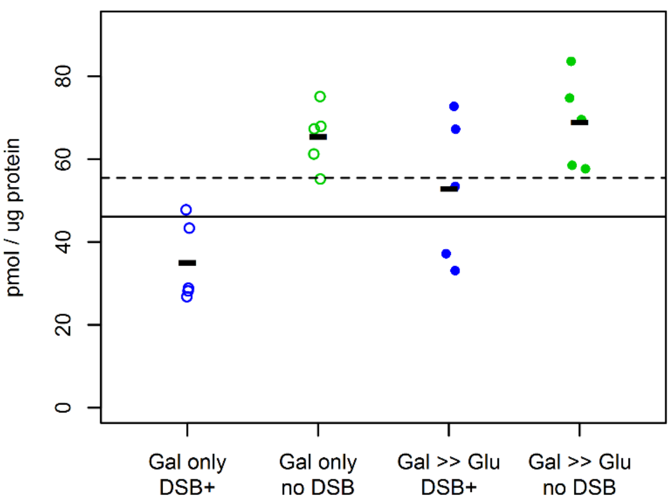

**Phosphoenolpyruvate**

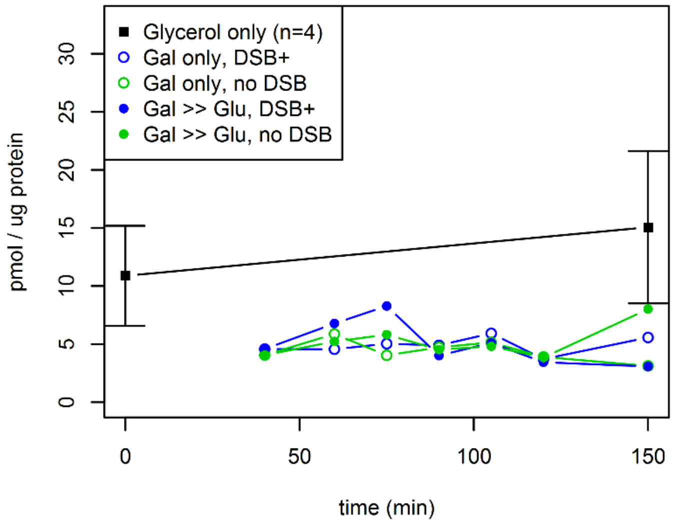


**Phosphoenolpyruvate**

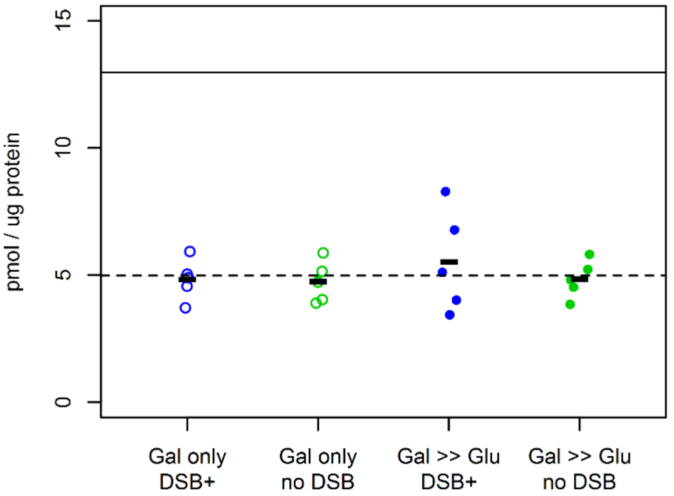


**Lactate**

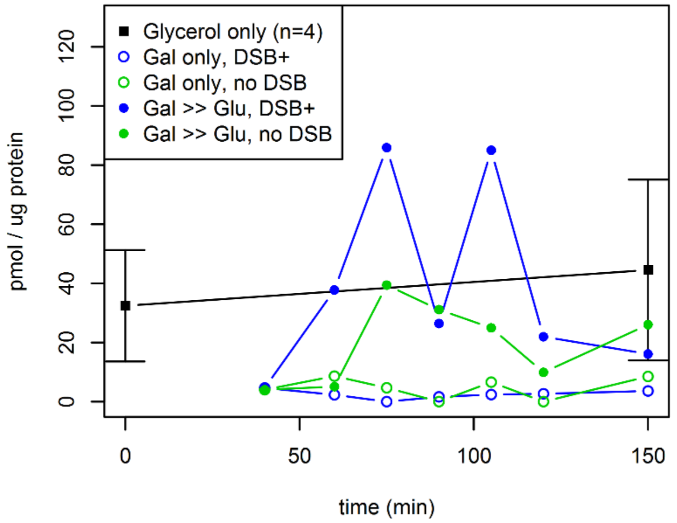


**Lactate**

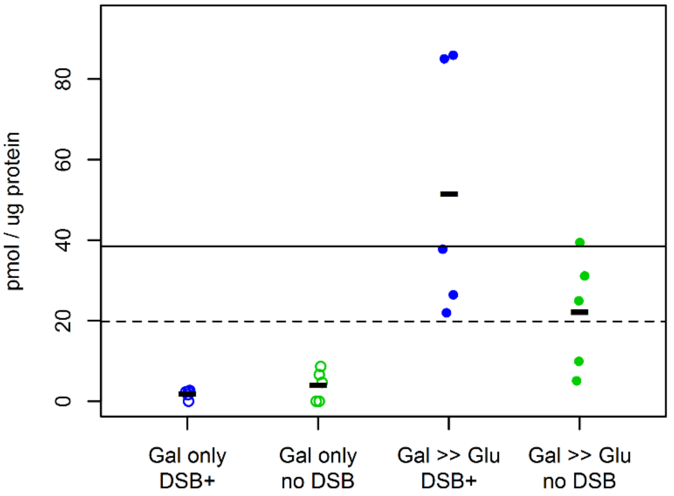


**Citrate + Isocitrate**

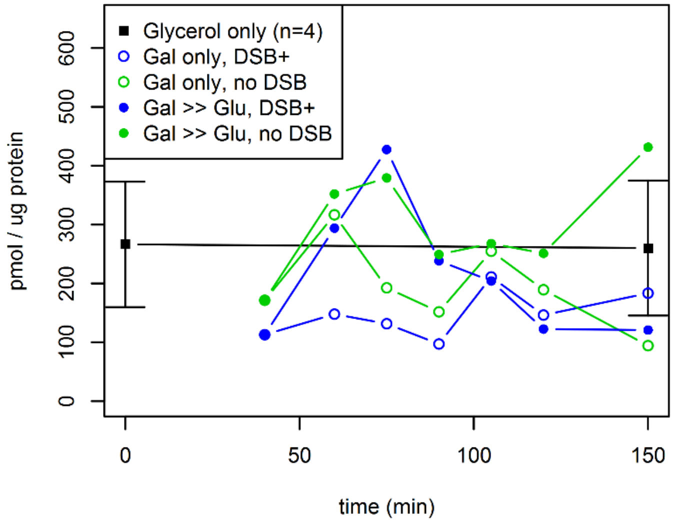


**Citrate + Isocitrate**

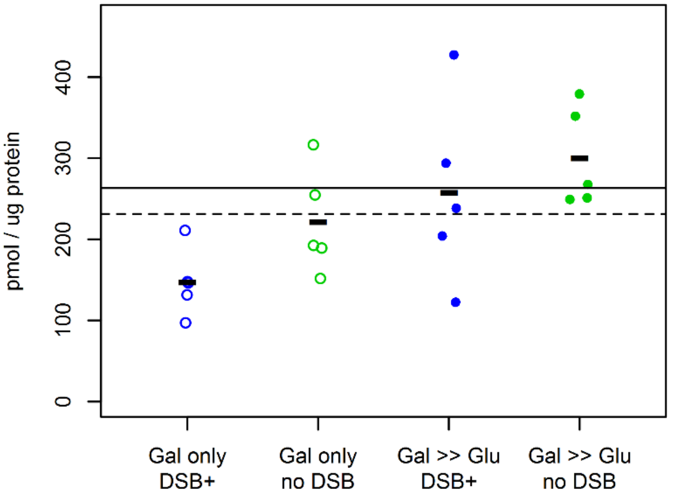

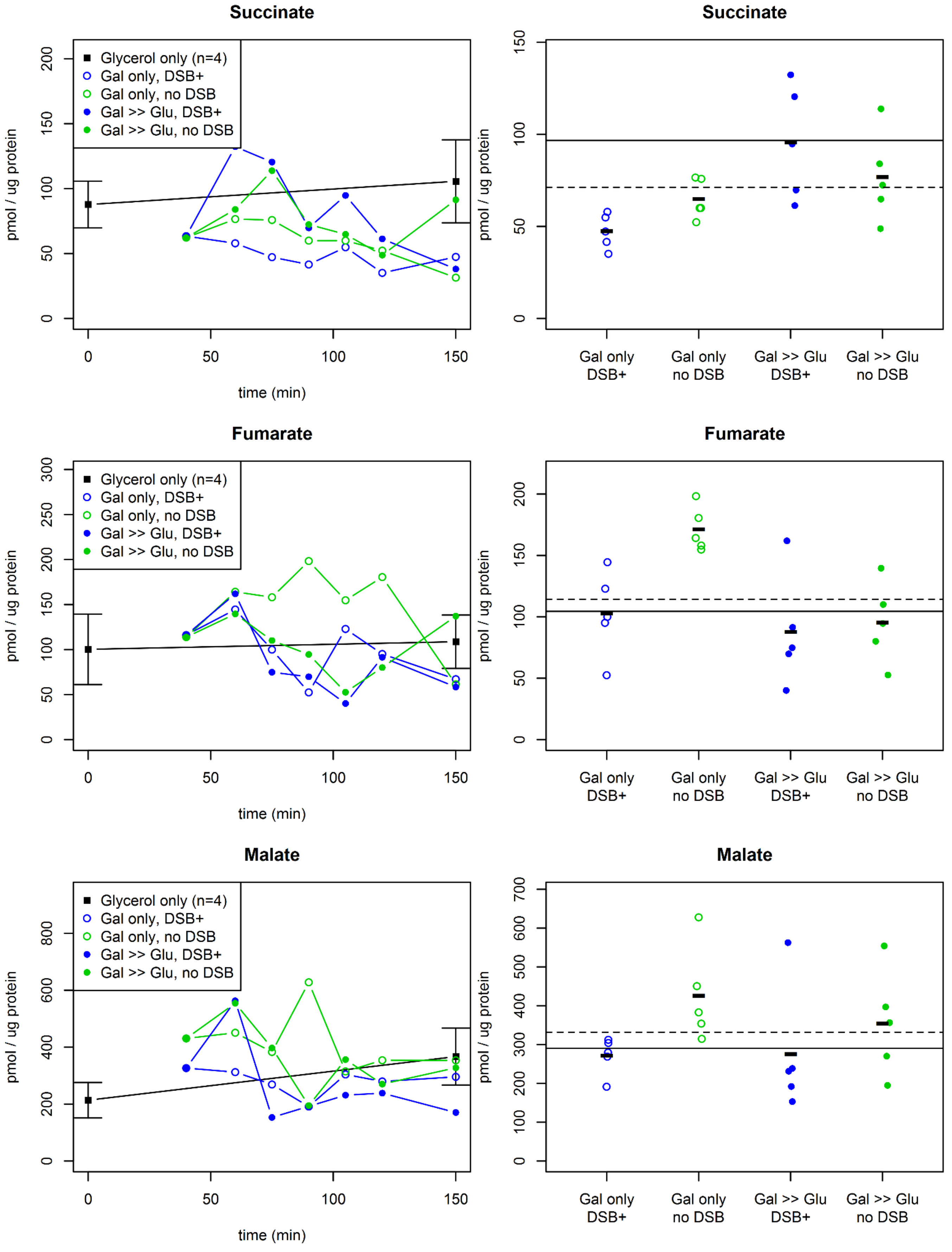
Succinate
Glycerol only (n=4)
Gal only, DSB+
Gal only, no DSB
Gal >> Glu, DSB+
Gal >> Glu, no DSB
pmol / ug protein
0
50
100
150
200
time (min)
Succinate
Gal only DSB+
Gal only no DSB
Gal >> Glu DSB+
Gal >> Glu no DSB
Fumarate
Glycerol only (n=4)
Gal only, DSB+
Gal only, no DSB
Gal >> Glu, DSB+
Gal >> Glu, no DSB
250
300
time (min)
Fumarate
Malate
Glycerol only (n=4)
Gal only, DSB+
Gal only, no DSB
Gal >> Glu, DSB+
Gal >> Glu, no DSB
400
600
800
time (min)
Malate
300
500
700

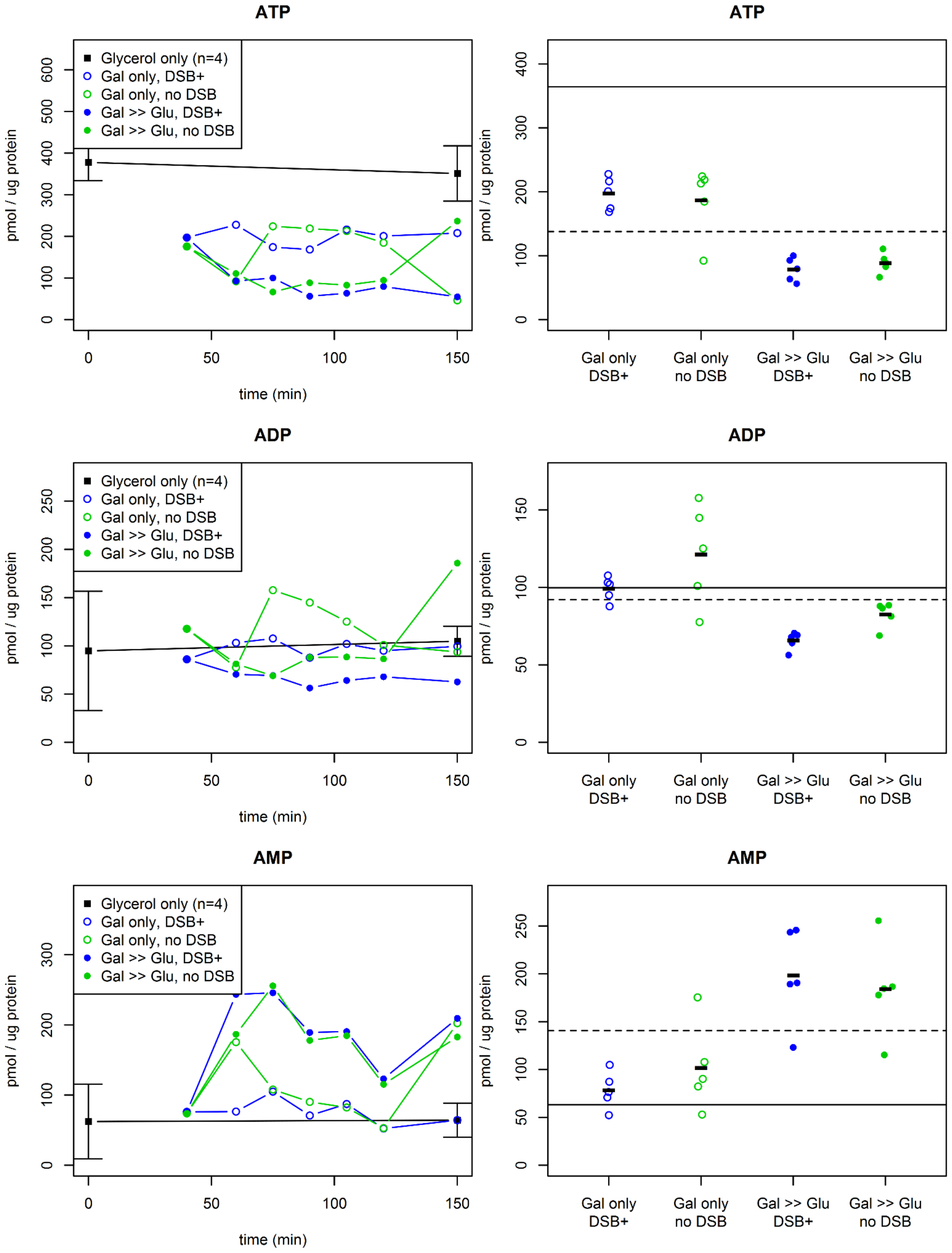
ATP
Glycerol only (n=4)
Gal only, DSB+
Gal only, no DSB
Gal >> Glu, DSB+
Gal >> Glu, no DSB
pmol / ug protein
time (min)
ATP
pmol / ug protein
Gal only DSB+
Gal only no DSB
Gal >> Glu DSB+
Gal >> Glu no DSB
ADP
Glycerol only (n=4)
Gal only, DSB+
Gal only, no DSB
Gal >> Glu, DSB+
Gal >> Glu, no DSB
pmol / ug protein
time (min)
ADP
pmol / ug protein
Gal only DSB+
Gal only no DSB
Gal >> Glu DSB+
Gal >> Glu no DSB
AMP
Glycerol only (n=4)
Gal only, DSB+
Gal only, no DSB
Gal >> Glu, DSB+
Gal >> Glu, no DSB
pmol / ug protein
time (min)
AMP
pmol / ug protein
Gal only DSB+
Gal only no DSB
Gal >> Glu DSB+
Gal >> Glu no DSB

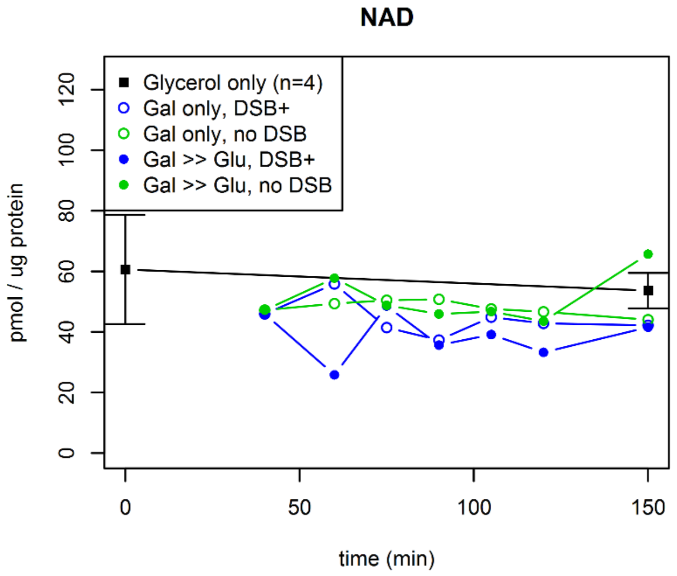
NAD
Glycerol only (n=4)
Gal only, DSB+
Gal only, no DSB
Gal >> Glu, DSB+
Gal >> Glu, no DSB
pmol / ug protein
0
20
40
60
80
100
120
0
50
100
150
time (min)

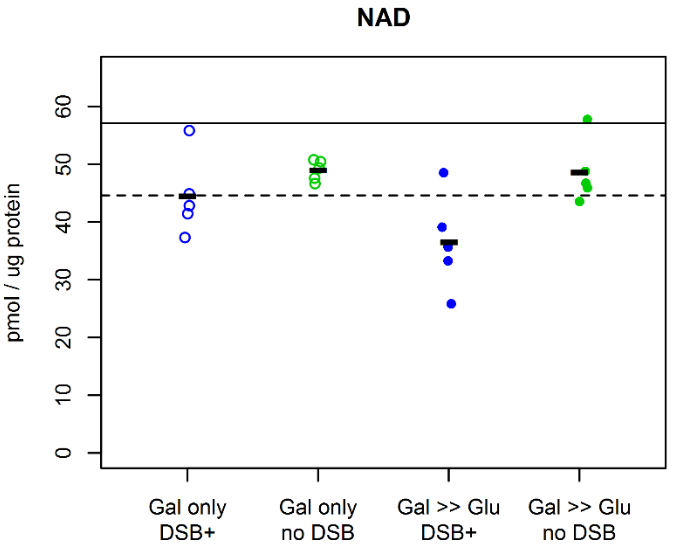
NAD
pmol / ug protein
0
10
20
30
40
50
60
Gal only
DSB+
Gal only
no DSB
Gal >> Glu
DSB+
Gal >> Glu
no DSB

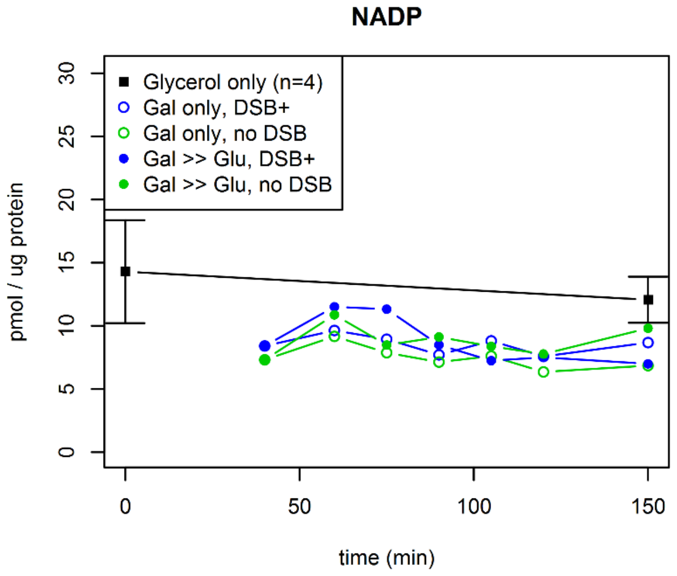
NADP
Glycerol only (n=4)
Gal only, DSB+
Gal only, no DSB
Gal >> Glu, DSB+
Gal >> Glu, no DSB
pmol / ug protein
0
5
10
15
20
25
30
0
50
100
150
time (min)

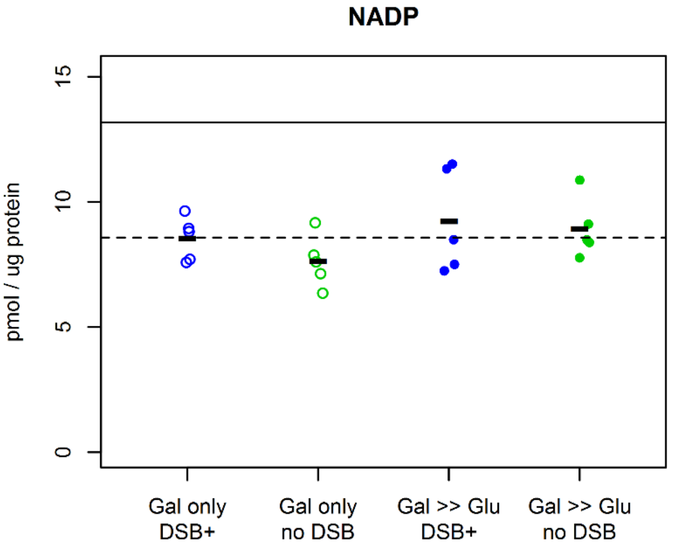
NADP
pmol / ug protein
0
5
10
15
Gal only
DSB+
Gal only
no DSB
Gal >> Glu
DSB+
Gal >> Glu
no DSB

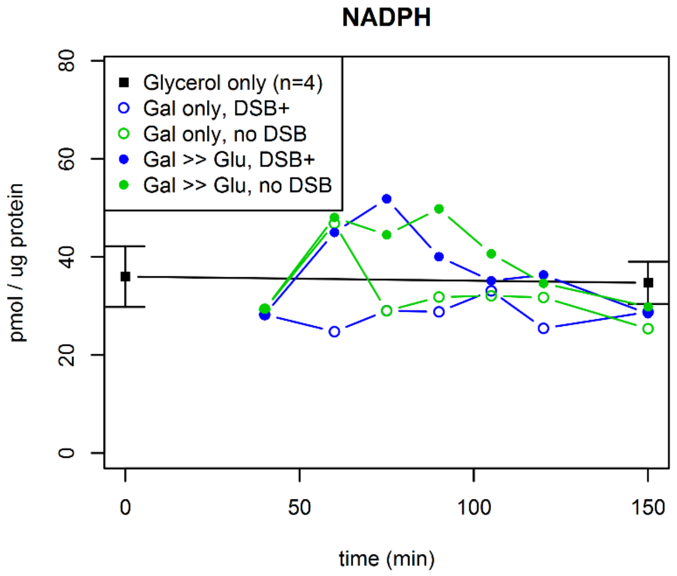
NADPH
Glycerol only (n=4)
Gal only, DSB+
Gal only, no DSB
Gal >> Glu, DSB+
Gal >> Glu, no DSB
pmol / ug protein
time (min)

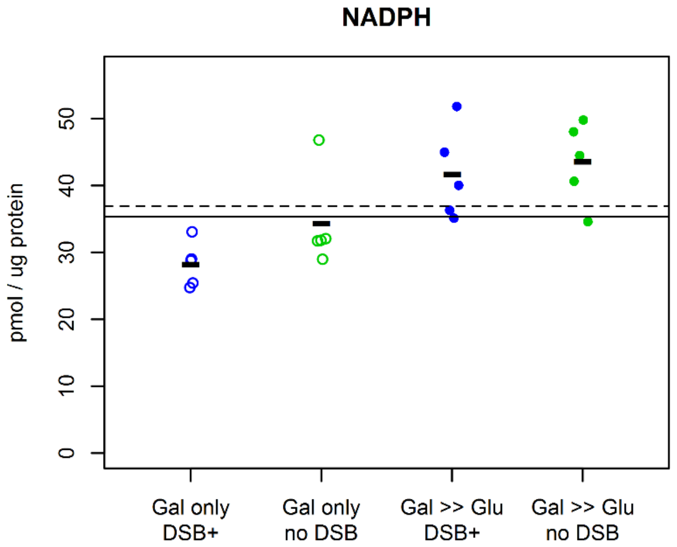
NADPH
pmol / ug protein
Gal only DSB+
Gal only no DSB
Gal >> Glu DSB+
Gal >> Glu no DSB

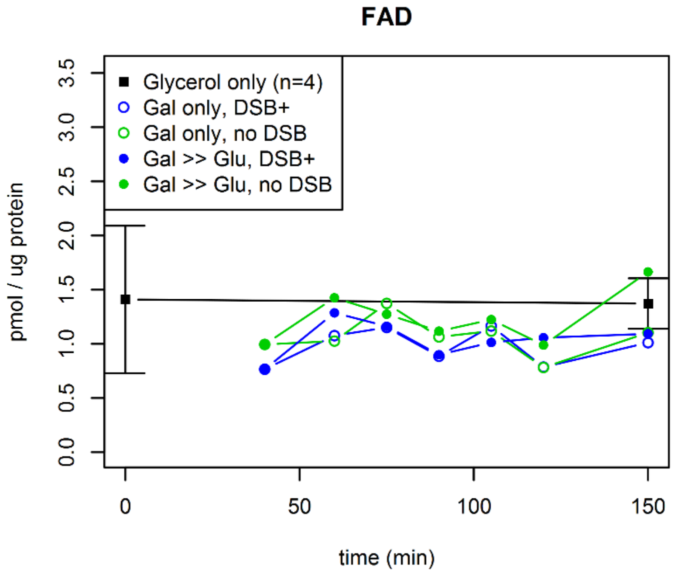
FAD
Glycerol only (n=4)
Gal only, DSB+
Gal only, no DSB
Gal >> Glu, DSB+
Gal >> Glu, no DSB
pmol / ug protein
time (min)

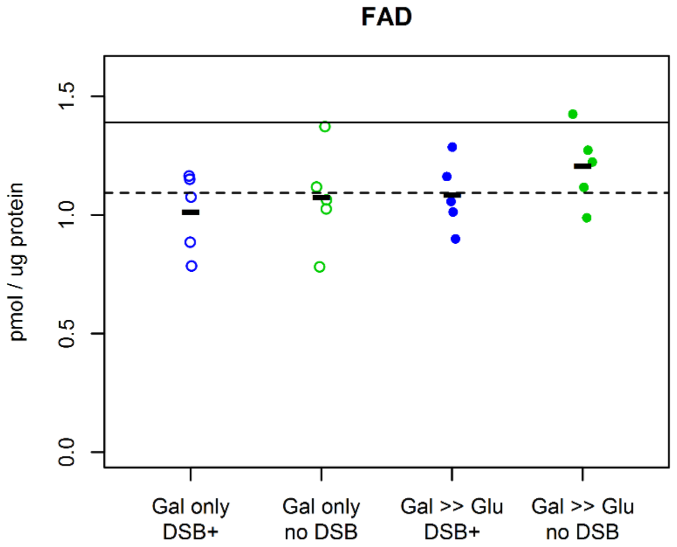
FAD
pmol / ug protein
Gal only DSB+
Gal only no DSB
Gal >> Glu DSB+
Gal >> Glu no DSB
